# Oncogenic KRAS engages an RSK1/NF1 complex in pancreatic cancer

**DOI:** 10.1101/2020.09.14.295394

**Authors:** Derek K. Cheng, Tobiloba E. Oni, Youngkyu Park, Jennifer S. Thalappillil, Hsiu-Chi Ting, Brinda Alagesan, Nadia Prasad, Keith D. Rivera, Darryl J. Pappin, Linda Van Aelst, David A. Tuveson

**Affiliations:** Stony Brook University; Cold Spring Harbor Laboratory

**Keywords:** KRAS, BioID, RSK, PDAC, NF1

## Abstract

Pancreatic ductal adenocarcinoma (PDAC) is a lethal malignancy with limited treatment options. Although activating mutations of the KRAS GTPase are the predominant dependency present in >90% of PDAC patients, targeting KRAS mutants directly has been challenging in PDAC.

Similarly, strategies targeting known KRAS downstream effectors have had limited clinical success due to feedback mechanisms, alternate pathways and toxicity due to the targeting of normal tissues. Therefore, identifying additional functionally relevant KRAS interactions in PDAC may allow for a better understanding of feedback mechanisms and unveil new potential therapeutic targets. Here, we used proximity labelling to identify protein interactors of active KRAS in PDAC cells. Fusions of wildtype (BirA-KRAS4B), mutant (BirA-KRAS4B^G12D^) and non-transforming and cytosolic double mutant (BirA-KRAS4B^G12D/C185S^) KRAS with the BirA biotin ligase were expressed in murine PDAC cells. Mass spectrometry analysis revealed that RSK1 was enriched among proteins that selectively interacted with membrane-bound KRAS^G12D^. RSK1 required the NF1 and SPRED proteins to interact with KRAS-GTP at the membrane. In both murine and human PDAC lines, membrane-targeted RSK1 was tolerated but inhibited cell proliferation following oncogenic KRAS abrogation to reveal a negative feedback role for membrane-localized RSK1 on wild-type KRAS. Inhibition of the wild-type KRAS, which has been previously proposed to suppress KRAS oncogenesis, may partially explain how RSK1 has been identified as a dependency in some KRAS mutant cells and may provide an additional function for NF1 in tumorigenesis.

**Significance Statement:** For decades, KRAS interactors have been sought after as potential therapeutic targets in KRAS mutant cancers, especially pancreatic adenocarcinoma (PDAC). Our proximity labeling screen with KRAS in PDAC cells highlight RSK1 as a notable mutant-specific interactor. Functionally, we show that RSK1 mediates negative feedback on wild-type KRAS in PDAC cells.

## Introduction

56,770 new cases of pancreatic cancer were estimated for 2020 and despite limited treatment options, the 5-year relative survival rate has consistently remained below 10% [1]. About 85% of these pancreatic cancer tumors are pancreatic ductal adenocarcinoma (PDAC) [2]. Poor outcomes of PDAC cases result from late diagnoses leading to unresectable and heterogeneous tumors as well as ineffective therapies, which only prolong survival on the order of months [3–5]. Over 90% of PDAC contain KRAS mutations and are associated with poor prognosis [6]. Furthermore, mice expressing mutant KRAS in the pancreas develop precursor lesions, which sporadically progress into PDAC, and this progression is potentiated when combined with other mutations or deletion of tumor suppressor genes [7–11]. Additionally, independent studies have shown that the maintenance of murine PDAC cells require KRAS[12–14].

KRAS as a Ras GTPase acts as a molecular switch at the plasma membrane that relays growth factor signaling from receptor tyrosine kinases to downstream pathways such as RAF/MEK and PI3K/AKT [15]. GTP binding alters the conformation of the KRAS G-domain, thereby creating binding sites for downstream effectors to trigger enzymatic cascades that promote cell transformation [16–19]. Intrinsically, KRAS slowly hydrolyzes GTP into GDP to halt signaling; however, GTPase activating proteins (GAPs) such as neurofibromin 1(NF1) catalyze this process [20]. In contrast, guanine nucleotide exchange factors (GEFs), such as SOS, catalyze the exchange of GTP for bound GDP. In most PDAC cases, KRAS is mutated at the 12^th^ residue located in the G-domain from glycine to either a valine (G12V) or more commonly, aspartate (G12D). These mutations sterically prevent the “arginine finger domain” of GAPs from entering the GTPase site, thereby blocking extrinsic allosteric GTPase activation and stabilizing RAS-GTP [21, 22]. Activating mutations in KRAS constitutively trigger RAF/MEK and PI3K/AKT pathways leading to increased cell proliferation as well as other pro-oncogenic behaviors [15]. KRAS signaling not only relies on the G-domain, but also the C-terminal hypervariable domain (HVR), which is required to stabilize KRAS on membranes where signaling is most efficient [23–26]. Independent studies suggest that specific biochemical and cellular consequences of KRAS activation are attributed to the unique properties of the HVR of the predominant splice form KRAS4B, namely the polybasic domain and the lipid anchor [27–30]. Localization of Ras proteins to the plasma membrane requires the prenylation of the CAAX motif [23]. Additionally, for KRAS4B, the hypervariable region contains a highly polybasic domain consisting of several consecutive lysines, which can interact with the negative charges on the polar heads of phospholipids and stabilize protein interactions [31]. Structural and biochemical characterization of the HVR and G domain has contributed to a better understanding of the signaling outputs of KRAS and led to KRAS-targeting strategies.

Various approaches to inhibit KRAS include direct inhibition, expression interference, mislocalization, and targeting of downstream effectors [32]. Thus far, direct inhibitors against KRAS have only successfully targeted the G12C mutant, which comprises 2.9% of KRAS mutant PDAC [21, 33]. For other KRAS mutants, targeting downstream effectors of KRAS in pancreatic cancer remains as an alternate approach. Unfortunately, dual inhibition of MEK and AKT pathways in pancreatic cancer patients resulted in increased adverse events compared to the standard second line chemotherapy, mFOLFOX, and did not improve overall survival [34]. Difficulty in targeting KRAS due to adaptive resistance and feedback regulation motivates a better understanding of KRAS biology [35]. For example, although PDAC typically features a mutant KRAS, there may be a role for its wild-type counterpart as well as the Ras paralogs, all of which are GAP sensitive and subject to signaling feedback. While oncogenic KRAS promotes the activation of wild type H- and N-RAS via allosteric stimulation of SOS1 [36], wild-type KRAS has considered as a tumor suppressor in some KRAS-mutant cancer contexts based on the observed mutant-specific allele imbalance that occurs throughout tumor progression. Additionally, the reintroduction of wild-type KRAS abolished tumor T-cell acute lymphoblastic leukemia development and impaired tumor growth in Kras mutant lung cancer cells *in vivo* [37–39]. While the ability of wild type KRAS to dimerize is essential for its tumor suppressive function, it is unclear whether it acheives this by preventing mutant homodimers or though unique wild type effectors [40]. The discovery of novel KRAS protein interactors involved in downstream signaling or feedback and compensatory pathways may elucidate why inhibition of downstream pathways have had limited clinical impact in PDAC. Here, we perform proximity labelling experiments by expressing a fusion of BirA^R118G^ biotin ligase and KRAS in PDAC cells which, in the presence of high concentrations of biotin, generates reactive biotinoyl-AMP that labels lysines of nearby proteins, such as interactors of its fusion partner KRAS [41–43]. The biotinylated interactor proteins can be isolated by streptavidin pulldown and analyzed by proteomics to identify novel protein interactors [44–46]. Because covalent labelling occurs in living cells, enzymatic labeling may potentially locate transient interactors and protein complexes.

Recently, several groups used BioID (proximity labelling method to identify KRAS interactors [47, 48]. While BirA-KRAS screens have been performed in 293T and colon cancer cells and have uncovered and validated the functional relevance of PIP5KA1 and mTORC2 in PDAC cells, a BirA-KRAS screen had yet to be directly performed in a PDAC model system prior to this study. The tumor context may determine protein expression and relevant interactions. Therefore, we sought to perform the first BirA-KRAS screen in PDAC cells. We hypothesize that proximity labeling with BioID presents a means for identifying new mutant KRAS-specific interactions in PDAC, which may unveil new insights into therapeutic design for this malignancy.

## Results

### BioID-KRAS in pancreatic ductal adenocarcinoma cells

To identify novel interactors of mutant KRAS in PDAC that helps better understand mutant KRAS biology and inform therapeutic efforts, we performed the BioID screening in mouse PDAC cells. To make the screen relevant to PDAC, our studies focused on using BioID with KRAS^G12D^, the most common mutation in PDAC, and performed the experiment in three cell lines derived from *FRT-LSL-Kras^G12V^-FRT; LSL-Tr<u>p</u>53^R172H^; Pdx1-<u>C</u>re;R26-FlpOERT2* (FPC) mouse PDAC tumors [49]. The endogenous expression of KRAS^G12V^ in this mouse model and isolated cell lines makes its detection by mass spectrometry (MS) distinguishable from our exogenously expressed BirA-Kras^G12D^. The endogenous *Kras^G12V^* allele can be excised upon 4-hydroxy-tamoxifen (4-OHT) treatment in these PDAC cells to minimize any competition from endogenous mutant KRAS that may confound the identification of BirA-KRAS interactors.

To label proteins proximal to KRAS in PDAC cells, we designed and expressed various BioID fusion proteins with BirA and Kras4B, the Kras splice isoform which is ubiquitiously expressed and upregulated in some cancer cells (Fig. 1A) [28]. We fused both mutant (*KRAS^G12D^*) and wildtype murine *Kras* (*KRAS^WT^*) to the C terminus of myc-tagged *BirA* to create *BirA-Kras^G12D^* (B-G12D) and *BirA-Kras^WT^* (B-WT). Since both the polybasic domain and the CAAX motif at the C-terminus are required for prenylation for membrane localization which is critical for KRAS function, we used a membrane-localized BirA control (B-CAAX) by fusing the last 20 amino acids of murine KRAS to BirA [50, 51]. Interactions with active KRAS that preferentially occur at the membrane may be more relevant to KRAS signaling [52, 53]. To determine whether any membrane-preferred interactions exist, we created a double mutant BirA-KRAS fusion protein including an additional mutation in the KRAS CAAX motif (B-G12D/C185S) as a non-transforming cytosolic comparator.

**Figure 1.**
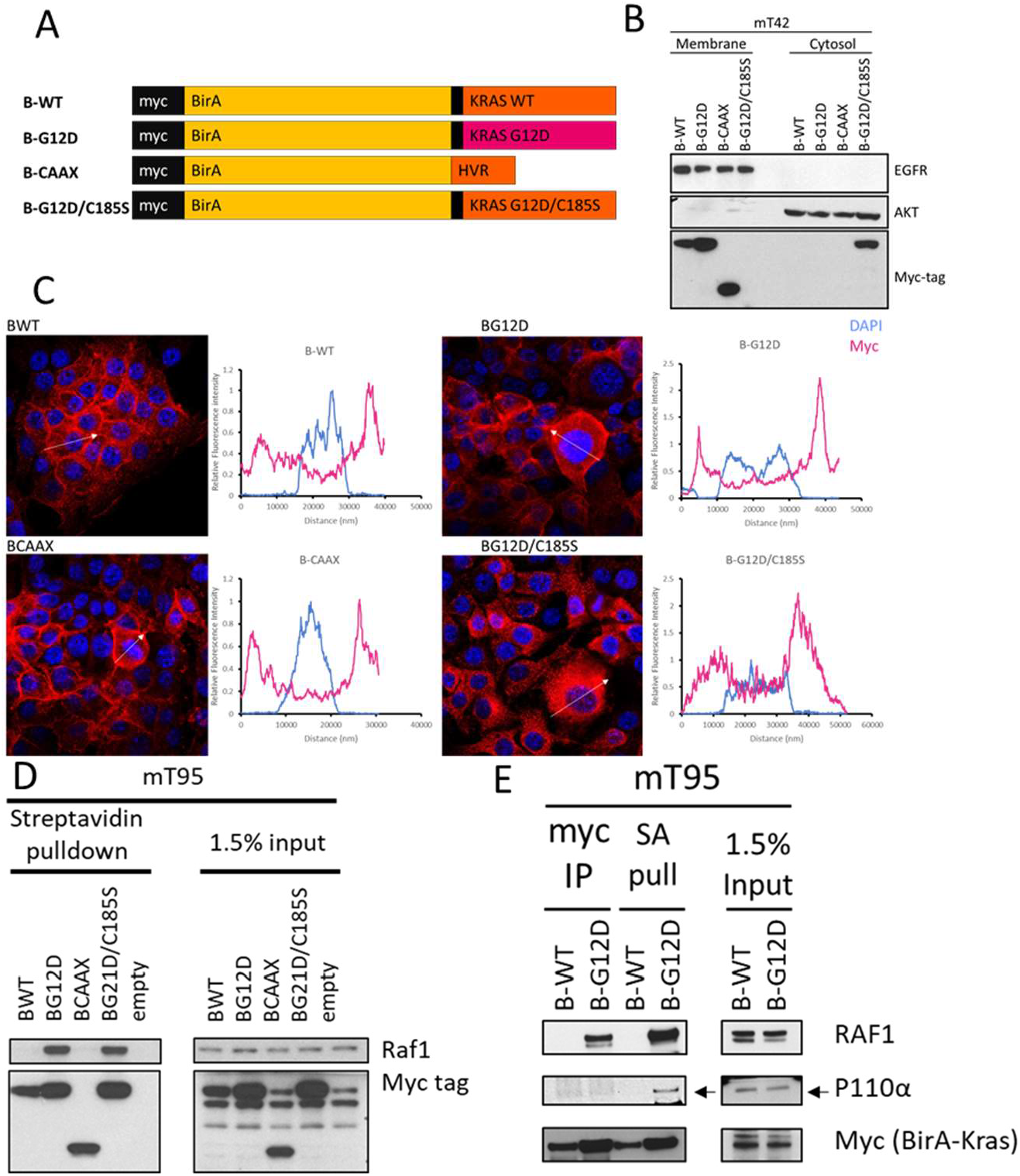
Properties of BirA-KRAS fusions. A) Design of BirA-KRAS fusion constructs. Myc-tagged BirA was fused to the N-terminus of murine wild-type KRAS4B and mutant KRAS4BG12D with a short glycine linker (GGSG). Membrane-localized BirA was created by fusing the last 20 amino acids of murine KRAS to myc tagged BirA. KRAS4BG12D/C185S was created by mutating the cysteine residue in the CAAX prenylation motif to mislocalize the protein. B) mT42-2D cells expressing B-WT, B-G12D, B-CAAX and B-G12D/C185S were sequentially fractionated into cytosolic and membrane isolations and probed for myc-tag, as well as EGFR and AKT representing membrane and cytosolic controls, respectively. C) immunofluorescence (IF) of B-WT, B-G12D, B-CAAX and B-G12D/C185S in mT93-2D cells. Staining of the myc-tagged BirA fusions are represented in red while DAPI-stained nuclei are represented in blue in the IF images. The arrows indicate the profiles taken for the relative fluorescent intensities from one edge to the other edge in cells (adjacent panels). D) mT95-2D FPC cells were infected to express B-WT, B-G12D, B-CAAX, B-G12D/C185S and empty control. Cells were incubated with 50 μM biotin for 24 hours prior to lysis and streptavidin pulldown. E) Lysate from mT95 expressing B-WT and B-G12D was used for both immunoprecipitation and streptavidin pulldown under similar conditions and probed for RAF1, p110α, and the myc-tagged Bir-Kras fusions.

We first validated the localization of our panel of B-WT, B-G12D, B-CAAX and B-G12D/C185S proteins in mT42 FPC cells by isolating membrane and cytoplasmic fractions from these cells by sequential fractionation [54]. As expected, B-WT, B-G12D and B-CAAX proteins localized to the membrane, while the CAAX mutant B-G12D/C185S localized to the cytoplasm (Fig 1B). These findings were supported by myc-tag immunofluorescence (IF) stains of B-WT, B-G12D, and B-CAAX that exhibited a plasma membrane profile, and predominantly cytosolic IF detection of B-G12D/C185S in PDAC cells (Fig 1C). These experiments confirm that the HVR of KRAS4B is essential and sufficient to localize fusion proteins to the plasma membrane [50, 51].

To ensure that the constructs were functional, we confirmed that B-G12D exclusively activated Ras pathways, MAPK and AKT, in 293T and 3T3 cells, respectively (Fig S1A, S1B), Additionally, among the BirA-KRAS fusion proteins, only B-G12D increased cell proliferation and conferred anchorage-independent growth. To determine the functionality of BirA, we then infected FPC cells to express the B-WT, B-G12D, B-CAAX and B-G12D/C185S fusion constructs and treated the cells with biotin to determine whether the known Ras interactor RAF1 would be preferentially biotinylated by any of these fusion proteins. Indeed, B-G12D preferentially biotinylated more Raf1 than B-WT and B-CAAX, but biotinylated comparable levels of RAF1 as B-G12D/C185S (Fig 1D). This finding suggests that the interaction of RAF1 with KRAS depends on the GTP-bound state of KRAS rather than its localization and does not necessarily cause Ras pathway activation.

Having characterized functional aspects of our BirA-KRAS fusion proteins, we sought to determine whether proximity labeling is more effective than immunoprecipitation to identify protein interactors of KRAS as this would dictate the technique used for exploratory MS experiments. To test this hypothesis, we harvested protein lysate from biotin-treated B-WT and B-G12D-expressing FPC cells for both a streptavidin pulldown and an immunoprecipitation followed by western blot analysis. Given the same input amount and conditions, both approaches were able to detect similar amounts of RAF1, whereas streptavidin pulldown was superior in detecting the p110α subunit of PI3-Kinase (Fig 1E). Stoichiometry may limit the immunoprecipitation approach such that only a fraction of enriched B-G12D molecules co-precipitates p110α; however, in the proximity labeling approach a single molecule of B-G12D may biotinylate several molecules of p110α, increasing its detection by streptavidin pulldown. Because BioID was more sensitive in detecting the interaction of p110α with KRAS than immunoprecipitation, we set out to employ this approach to identify new interactors by MS.

### Identification of membrane- and mutant-specific interactors of oncogenic KRAS

To detect novel interactors of oncogenic KRAS, we performed BioID-MS analysis on 3 biological replicates of murine FPC tumor cells (mT42, mT93, mT95). For each cell line, we generated 4 lines stably expressing each of our BioID constructs (B-WT, B-G12D, B-CAAX, B-G12D/C185S) to detect membrane-specific and/or mutant specific interactors by comparing B-G12D to B-WT and/or B-G12D to B-G12D/C185S respectively (Fig. 2A). To learn whether endogenous mutant Kras^G12V^ could affect the interactors identified, these cells were treated with 4-OHT to ablate the expression of Kras^G12V^ or DMSO as a vehicle control for 3 days prior to biotin treatment, followed by lysate harvest, streptavidin pulldown and mass spectrometry analysis. To account for differences of BirA expression in each cell line and the variability in the number of peptides in each run for MS, we normalized the quantitative peptide abundances to the BirA counts for each experiment.

**Figure 2.**
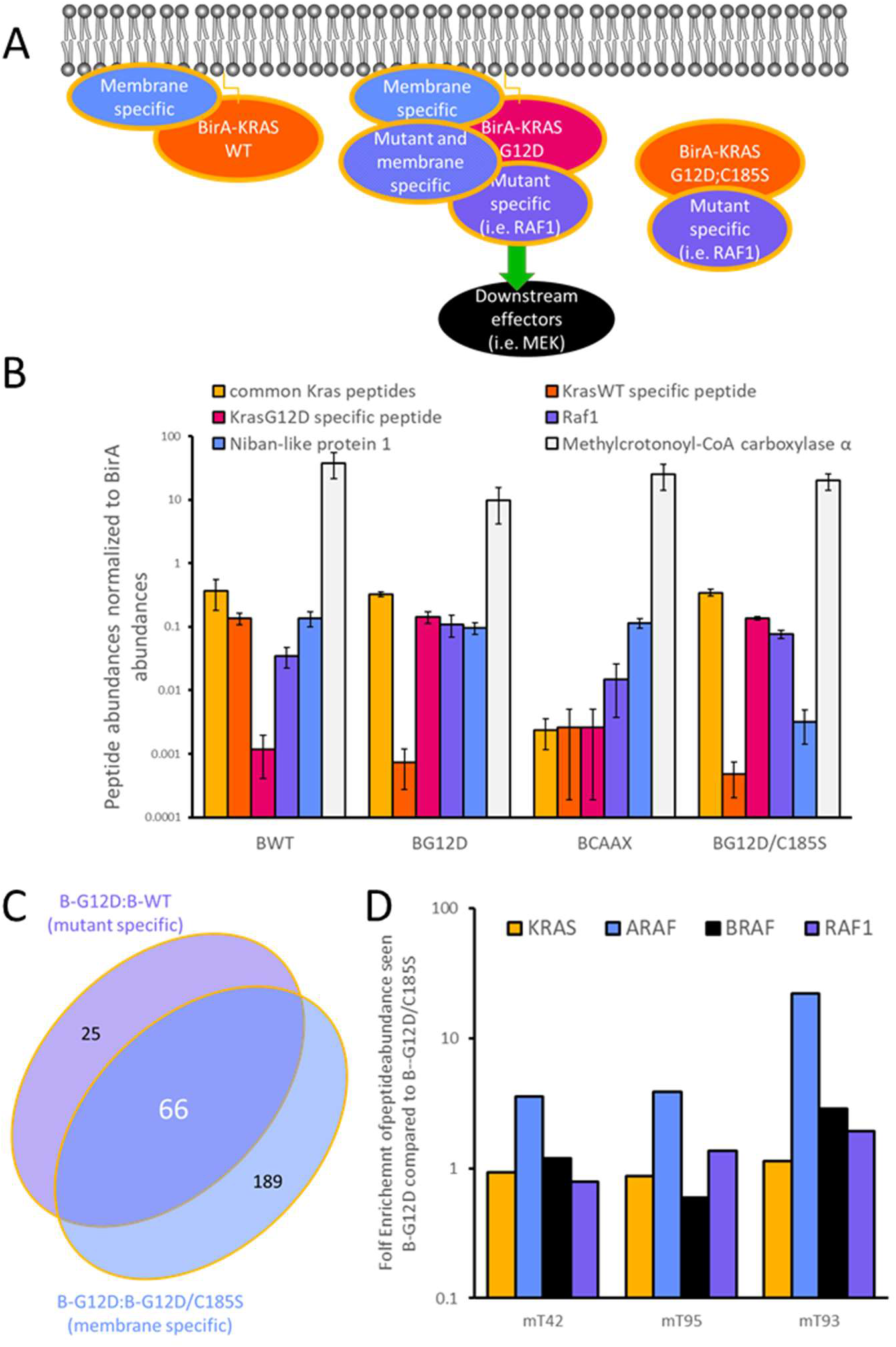
BioID-Kras identifies mutant and membrane specific interactors. A) Schematic of determining mutant-specific interaction by comparing B-G12D to B-WT and membranespecific interactions by comparing B-G12D to B-G12D/C185S. B) Mass spectrometry relative quantification of internal control KRAS peptides and positive controls (RAF1 and niban-like 1) after normalization to BirA counts for B-WT, B-G12D, B-CAAX and B-G12D/C185S samples from vehicle-treated FPC cells. Error bars represent standard error of the mean (n=3). C) Venn diagram of overlap between protein candidates enriched in B-G12D compared to B-WT indicating mutant specificity as well as compared to B-G12D/C185S indicating membrane specificity. D) Enrichment of Raf family member peptides and KRAS peptides in B-G12D samples compared to B-G12D/C185S samples after BirA normalization.

Several observed peptides validated this approach for both the vehicle-treated (Fig 2B) and 4-OHT treated cells (Fig S2). MS detected shared KRAS peptides and also identified the G12-containing peptide across the three full length BirA-KRAS variants. Interestingly, G12V peptides were not found in any samples run suggesting that either the BirA tag prevents dimer formation or that heterodimers may be too scarce to detect. Additionally, samples with an intact CAAX box enriched for membrane-localized proteins, such as niban-like protein 1 (FAM129B), compared to samples from cytosolic B-G12D/C185S. Consistent with our western blot results (seen in Fig 1D), B-G12D and B-G12D/C185S had similar biotinylation levels of Raf1, whereas B-WT and B-CAAX labeled less RAF1.

To identify novel Kras^G12D^ effectors, we compared the proteins enriched by B-G12D and B-WT KRAS from our BioID-MS experiments and nominated 91 protein matches that met the criteria of being enriched in at least 2 out 3 biological replicates in the setting of an endogenous mutant KRAS^G12V^ allele. To further prioritize this list, we queried for membrane-specific candidates. Of the initial list of 91 mutant-specific candidate interactors, 66 proteins met the criteria of being enriched by B-G12D compared to B-G12D/C185S in at least 2 out 3 biological replicates (Table S1, Fig 2C). Interestingly, when sorting by membrane enrichment, ARAF was preferentially biotinylated at the membrane whereas RAF1 and BRAF were both comparably labelled by B-G12D and B-G12D/C185S (Table S2, Fig 2D). Although the direct interaction of KRAS and ARAF is well-established, these data suggest that our approach could map spatially distinct proximal interactors and uncover dynamic Kras^G12D^ interactors on the membrane.

To further refine our list and prevent confounding effects of endogenous mutant KRAS due to competition for substrates, we also performed BioID-MS analysis on the same BirA-KRAS expressing FPC cells treated with 4-OHT to remove the endogenous mutant KRAS^G12V^ allele (Fig 3A, S3A). By prioritizing candidates that were nominated in the mutant-specific and membrane-specific comparison for both the vehicle and 4-OHT conditions, we further narrowed our list to 32 proteins (Fig 3B). 11 of these protein matches were known interactors or KRAS effectors, including RAF family members (ARAF, BRAF, RAF1), SPRED1, SPRED2, NF1, MLLT4, RIN1, and mTORC2 complex members (MAPKAP1, RICTOR, DEPTOR) (Figure 3C, Table S3). Of the remaining 21 candidates, PLEKHA2, MYO1E and RPS6KA1 (RSK1) were also enriched in the B-G12D to B-WT comparison as previously reported in a BioID screen in 293 cell lines and RSK1 was enriched in a BirA-KRAS^Q61H^ to BirA-CAAX comparison of a 293 cell mapping project [48, 55]. Several previously reported Ras effectors including PI3K, KSR, RALBP, SOS1 and PIP5KA1 were not detected in our MS data set, suggesting that additional approaches may be need to increase the sensitivity of BioID assay in PDAC cells. Nonetheless, ARAF followed by RSK1 were the most highly ranked as consistently B-G12D enriched over B-WT in the 4-OHT setting.

**Figure 3.**
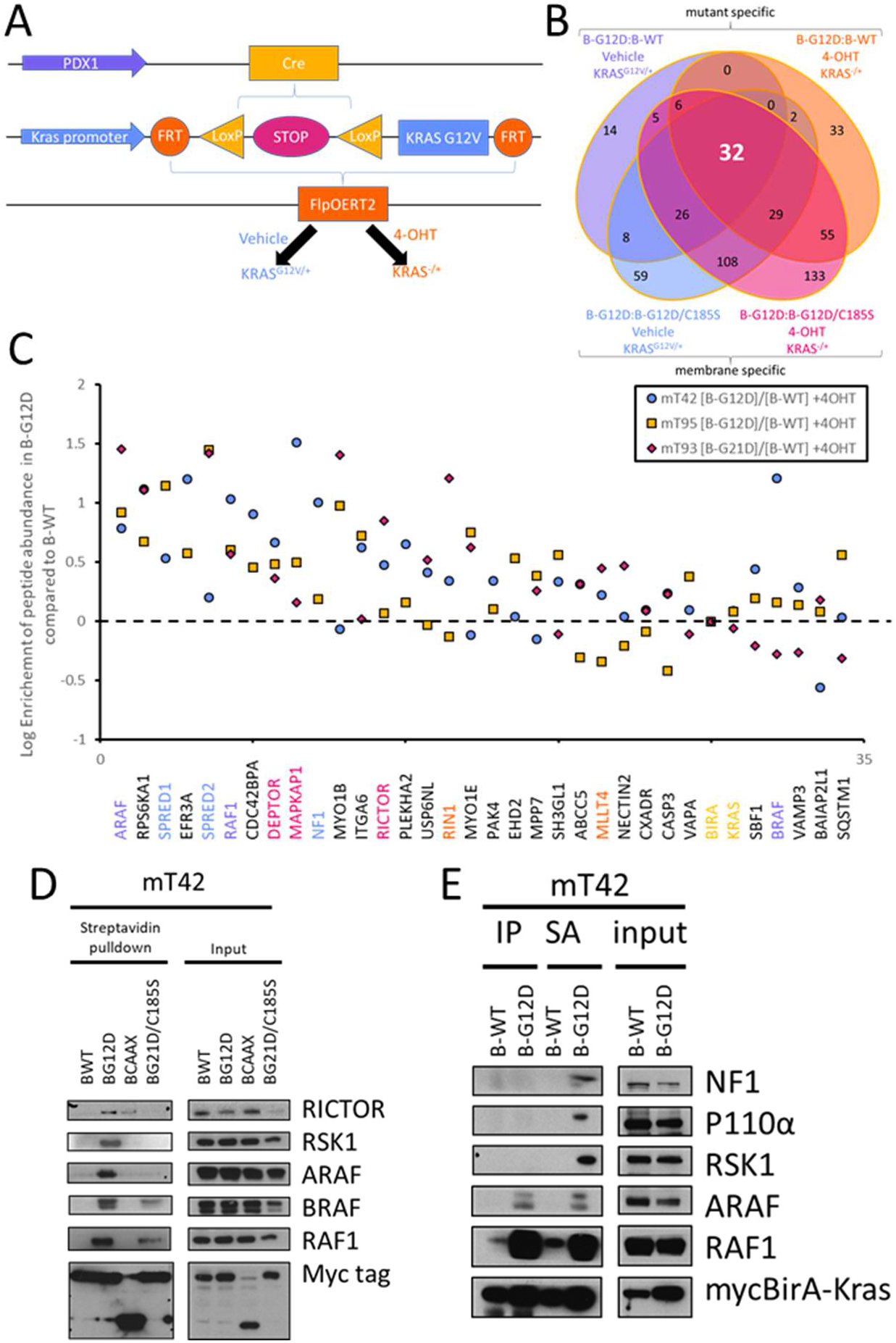
Proximity labelling nominates RSK1 as a mutant- and membrane-specific KRAS interactor. A) FPC mouse model features FRT recombinase sites flanking the *LSL-Kras^G12V^* allele use to remove the endogenously expressed KRAS^G12V^ B) Venn diagram of overlap of candidates that meet criteria for being mutant-specific and membrane-specific in both the 4-OHT treated and control treated conditions for 2 out of 3 biological replicates. 32 candidates met all criteria. C) Enrichment of peptide abundances for the 32 candidate interactors as well as KRAS and BirA control peptides. Known KRAS interactors include RAF members (purple), NF1/SPRED proteins (blue), mTORC2 (magenta), and RBD containing proteins (orange) with the internal controls, BirA and KRAS, in yellow. D) Western blot confirmation of biotinylation of RICTOR, RSK1, ARAF, BRAF, and RAF1 as B-G12D substrates in mT42. E) immunoprecipitation and streptavidin pulldowns of myc-tag B-WT and B-G12D for the detection of NF1, p110α, RSK1, ARAF, RAF1, myc-tag.

Of the interactors that were nominated to be both KRAS^G12D^-specific and plasma membrane-localized, RICTOR of mTORC2, ARAF, and RSK1 were among those validated by western blot analysis of the BioID samples (Fig 3D, S3B). Unlike RAFs and the mTORC2 complex, RSK1 lacks a known Ras-binding domain (RBD), and the mechanism by which RSK1 approaches KRAS has not been determined. We repeated myc-tag coimmunoprecipitation alongside streptavidin pulldowns of lysates from FPC cells comparing B-WT and B-G12D to validate the RSK1-KRAS interaction. However, like other Ras interactors aside from Raf family members such as NF1 and p110α, RSK1 was only detected by streptavidin pulldown of B-G12D samples and failed to appreciably enrich by immunoprecipitation, motivating alternative approaches to demonstrate protein interaction (Fig 3E, S3C).

### MAPK activation dependent-RSK1/NF1/SPRED complex as a novel KRAS interactor

We reasoned that RSK1’s proximity to KRAS may depend on another protein in murine PDAC cells similar to the dependency of RICTOR on MAPKAP1 for proximity to KRAS [48]. To test this hypothesis, we used a combination of BioID and CRISPR/Cas9 KO technology. BioID enables the detection of transient or weak RSK1 interactors, which are difficult to identify by immunoprecipitation. Whereas, CRISPR-mediated ablation of specific proteins in the biotinylated interactome enables the identification of proteins that promote the proximity of RSK1 to KRAS^G12D^ on the membrane. We reasoned that mediators of the RSK1-KRAS^G12D^ interaction would co-enrich with RSK1 in our previous BioID-MS analysis. Therefore, we sought to determine whether knocking out some membrane-specific proximal interactors of B-G12D, including MAPKAP1, RICTOR and NF1, would abolish the proximity biotinylation of RSK1. To this end, we designed gRNAs against the genes encoding these proteins and stably expressed the guides in Cas9-expressing mT42 and mT93 cells. By depleting these proteins, we confirmed the dependence of RICTOR on MAPKAP1 to interact with KRAS; however, depletion of neither mTORC2 component decreased RSK1 biotinylation (Fig 4A). Interestingly, the biotinylation of RSK1 markedly decreased with the ablation of NF1 (Fig 4B). NF1 is known to interact with SPRED proteins [56, 57]. Accordingly, we tested guides against the two SPRED proteins found in our MS screen. Guides directed against SPRED1 and SPRED2 slightly decreased biotinylation of NF1 [56] (Fig 4C, S4), and guides targeting SPRED2 decreased the biotinylation of RSK1. Notably, RSK1, SPRED2, and NF1 were nominated as the top 13^th^, 14^th^ and 20^th^ interactors specific to KRAS^Q61H^ in the human cell map dataset [55]. Taken together, our results corroborate previous reports and indicate that the proximity of RSK1 to Kras^G12D^ is dependent on NF1, and RSK1 may directly interact with NF1.

**Figure 4.**
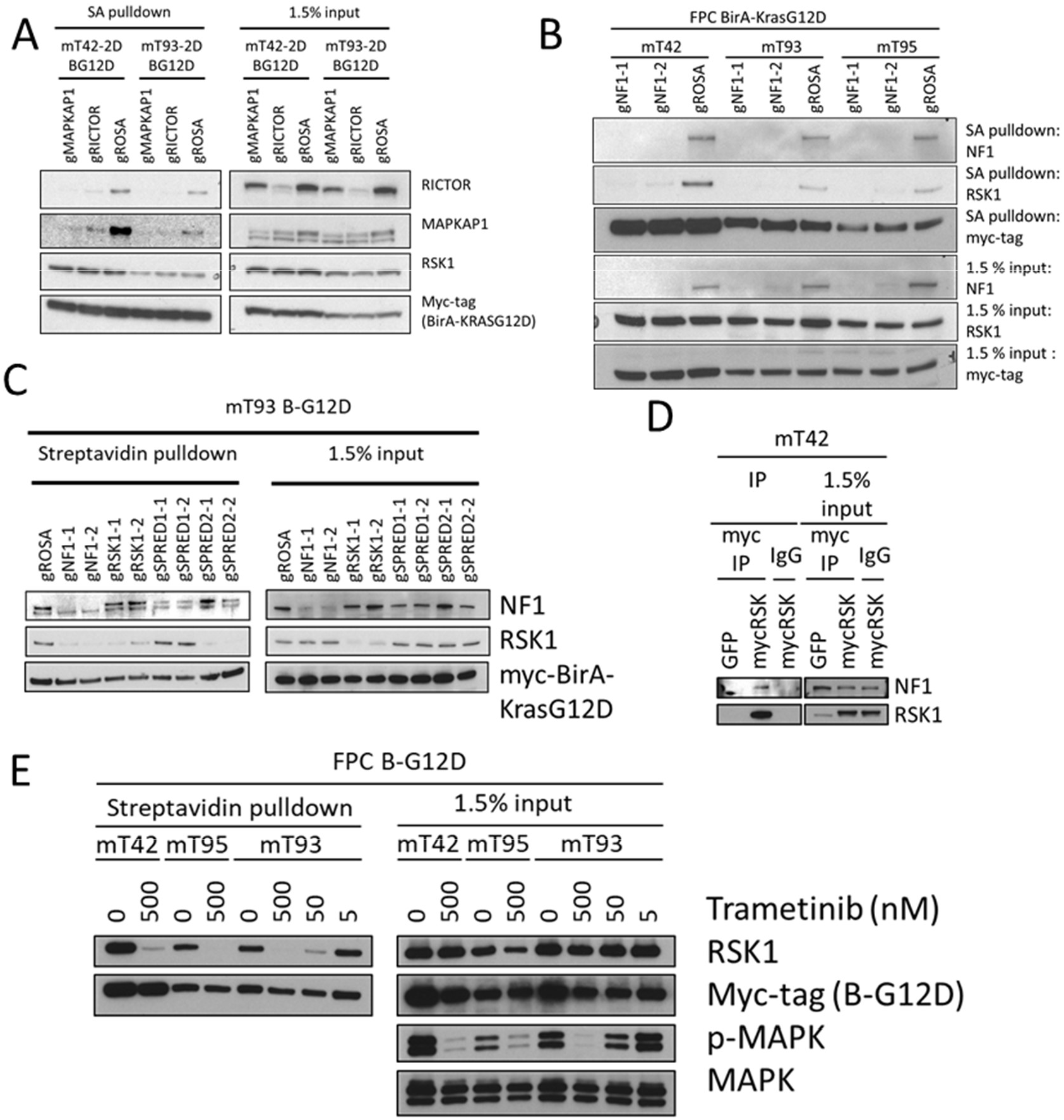
RSK1 is dependent on NF1 and MAPK activation to interact with KRAS. A) BioID of mT42 expressing B-G12D with CRISPR depletion of MAPKAP1 and RICTOR. B) BioID of mT42, mT93, and mT95 expressing B-G12D with CRISPR depletion of NF1. C) BioID of mT93 expressing B-G12D with CRISPR depletion of NF1, RSK1, SPRED1, and SPRED2. D) Myc-tag IP was performed on myc-tagged RSK1-expressing mT42 cells with GFP-expressing cells as the control. E) FPC cell lines expressing B-G12D were treated with trametinib (500 nM, 50 nM and 5 nM) during biotin incubation. The lysates and streptavidin pulldown were analyzed for the presence of RSK1, RSK2, NF1, ARAF and myc-tag as well as the phosphorylation status of MAPK.

In line with the putative functional interaction between NF1 and RSK1, the iGPS phosphorylation computational tool predicts NF1 to be a RSK1 substrate, and RSK1 has been shown to mediate negative feedback via NF1 [58, 59]. To validate the RSK1/NF1 interaction, we performed myc-tag co-immunoprecipitation in mT42 FPC cells expressing a myc-tagged RSK. Immunoblotting showed that the IP co-enriched myc-tagged Rsk1 and Nf1, providing additional evidence for a RSK1-NF1 complex as previously suggested by BioID (Fig 4D).

RSK1-4 are a family of serine/threonine kinases known to be activated by MAPK signaling and translocated to the plasma membrane transiently and in a MAPK-dependent manner upon EGF stimulation [60, 61]. To test whether the RSK1-KRAS interaction results from MEK/MAPK pathway activation, we treated B-G12D-expressing FPC cells with 500 nM of the MEK inhibitor, trametinib or vehicle control during the biotin incubation of the BioID assay. RSK1 biotinylation by B-G12D correlated with MAPK phosphorylation (Fig 4E), suggesting that the RSK1-KRAS interaction is dependent on MEK/MAPK pathway activation.

Having established a protein interaction complex of active KRAS that includes NF1 and RSK1, we next sought to determine a functional role of RSK1 in PDAC cells. RSK1 is a known effector downstream of the MAPK pathway and membrane-targeted RSK1, considered to be constitutively active, can confer MEK independence and mediate pro-survival signals in IL-3-dependent cells [62]. However, in other cellular contexts, membrane-localized RSK1 has also been shown to inhibit Ras signaling [63]. As KRAS localizes to the plasma membrane, we used a myristoylation (myr) tag to target RSK1 to the plasma membrane, confirmed by cell fractionation, to determine its function in relation to RAS [64] (Fig S5A). To identify whether membrane-targeted RSK1 negatively regulates the Ras pathway or contributes to Ras-mediated transformation in PDAC cells, we retrovirally introduced RSK1 and myr-tagged RSK1 into mT42 cells stably expressing our tagged KRAS variants, B-WT, B-G12D, and B-G12D/C185S. For comparison, we also included overexpression of ARAF and myr-tagged ARAF as well as an empty vector control. These cells were plated sparsely and treated with 4-OHT for 3 days to excise endogenous *Kras^G12V^* before crystal violet staining to measure foci formation (Fig 5A, S5B). mT42 cells expressing B-G12D formed colonies normally despite the removal of endogenous *Kras^G12V^*, whereas cells with overexpression of B-WT and B-G12D/C185S were unable to rescue this loss and exhibited less colonies. myr-tagged ARAF, but not untagged ARAF, was able to partially rescue the loss of Kras^G12V^ in both cells expressing B-WT and B-G12D/C185S, consistent with the observation that membrane-targeted ARAF does not require active KRAS to activate downstream signaling [65]. On the other hand, myr-RSK1 further inhibited colony formation of mT42 with either B-WT or B-G12D/C185S following excision of *Kras^G12V^*. While the mechanism of the inhibitory effect of membrane targeted RSK1 remains unknown, this effect is only seen in cells lacking active KRAS, suggesting that membrane-targeted RSK1 is playing a role upstream of KRAS.

**Figure 5.**
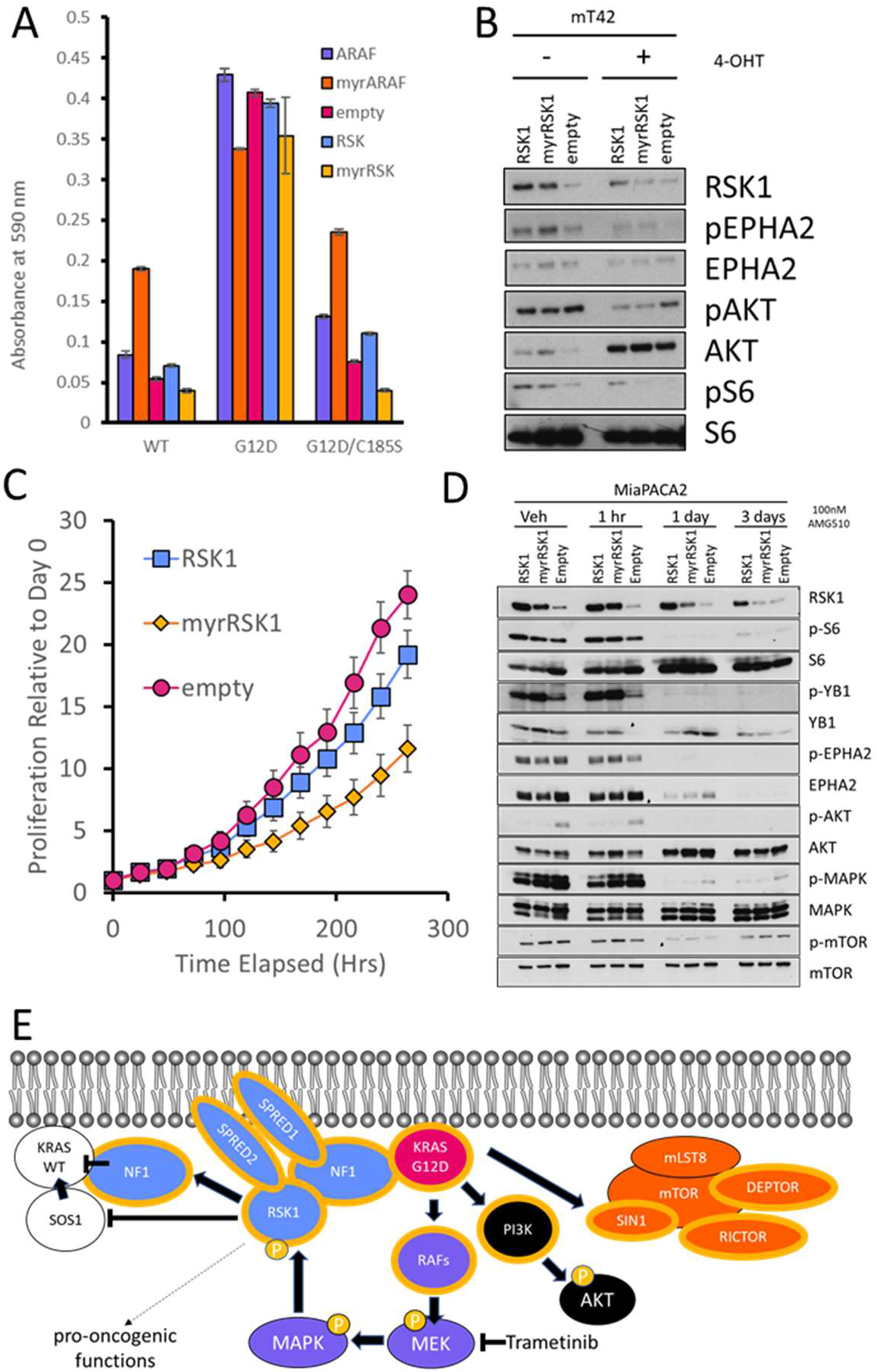
The role of RSK1 in a mutant KRAS-abrogated PDAC cell. A) mT42 cells expressing B-WT, B-G12D, B-G12D/C185S were infected to express ARAF, myr-tagged ARAF, RSK1, myr-tagged RSK1, and empty control. Post-selection, the cells were sparsely plated with 4-OHT and cultured for 3 additional days. The cells were stained with crystal violet, and bound dye extracted for quantification. Error bars indicate standard error of the mean (n=5). B). FPC cell line mT42 cells ectopically expressing RSK1, myr-RSK1, or GFP (empty) were treated with DMSO or 4-OHT, and immunoblotting was performed for RSK and related pathways. C) MiaPACA2 cells expressing RSK1 and myr-tagged RSK1, along with empty control, were treated with 100 nM of AMG510 for 1 hour, 1 day, and 3 days or DMSO control. Error bars represent standard error of the mean (n=. D) MiaPACA2 cells expressing RSK1, myr-RSK1, and GFP treated with 200 nM AMG510 were monitored for growth over time. E) Working model of the RAS/MAPK/RSK/NF1 feedback loop and effector pathways regarding proteins and complexes found to be biotinylated in BioID. Proteins outlined in yellow were preferentially biotinylated by B-G12D.

To confirm whether the inhibitory effect was due to inhibition of a Ras pathway biochemically, we introduced RSK1 and myr-RSK1 to mT42 cells and treated the cells with either 4-OHT or DMSO control. Immunoblotting revealed that RSK1 and myr-tagged RSK1 could be overexpressed in mT42 and these proteins were functional in increasing the phosphorylation of RSK1 substrates such as S6 and EPHA2 (Fig 5B). Upon KRAS abrogation, myr-RSK overexpression became particularly unstable compared to the untagged RSK1 overexpression. Additionally, AKT phosphorylation decreased in both cells that were exogenously expressing RSK and myrRSK, consistent with negative feedback on Ras signaling.

To extend these findings to human PDAC, RSK1 and myrRSK1 were ectopically expressed in MiaPACA2 cells, which harbor a Kras^G12C^ mutation that can be targeted by the commercially available KRAS^G12C^-specific inhibitor AMG510 [66]. To test whether membrane-localized RSK1 might affect cell proliferation as a function of the observed changes in RAS signaling pathways, we cultured RSK1, myrRSK1 and GFP (control) expressing MiaPACA2 cells with serial dilutions of AMG510 (5 μM, 1 μM, 200 nM, 40nM, 8nM, 1.6nM) (Fig S5C). While overexpression of untagged RSK1 did not affect MiaPACA2 proliferation compared to GFP control, the membrane-localized RSK1 sensitized cells to AMG510 treatment. At the 200nM AMG510 dosage, the growth of myr-RSK1 is diminished, consistent with a role of membrane-bound RSK1 as a negative regulator of wild-type RAS signaling (Fig 5C, S5C). To investigate the biochemical mechanism, we treated RSK1 and myr-RSK1 expressing cells with either 100 nM of AMG510 or DMSO control over a time course and performed an immunoblot of cell lysates to probe for RSK1 and the phosphorylation of RAS downstream effectors (MAPK and AKT) and RSK1 substrates (EPHA2 and YB-1) (Fig 5D, S5D [67, 68]. While the expression of RSK1 and myr-tagged RSK1 were stable under vehicle control treatment, administration of AMG 510 resulted in a decrease in myr-tagged RSK1 expression while the untagged RSK1 remained relatively constant. This finding suggests that cells with high expression of RSK1 at the membrane are not fit when dependent on wild-type Ras signaling. In the absence of AMG510, the phosphorylation of EPHA2 and YB-1 correlated with the expression of RSK1 regardless of myr-tag localization, confirming RSK1 function. Parallel to our findings in mT42 cells, MiaPACA2 cells showed less activation of MAPK in lysates with the highest levels of RSK1 under the same conditions. This trend also applied to the phosphorylation of AKT, another Ras effector. Treatment with AMG510 resulted in a significant decrease in phosphorylated AKT and MAPK by day 1. AMG510 decreasing the phosphorylation of MAPK and YB-1 can be explained by MAPK and YB-1 being both downstream of Ras and MEK with YB-1 being further downstream of MAPK and RSK1 [67–69]. Additionally, AMG510 decreased the expression of EPHA2 which is transcriptionally regulated by MAPK [68, 69].

These findings nominate RSK1 as an interactor of mutant KRAS that bridges the downstream activation of MAPK signaling to the upstream negative regulator of wild-type Ras, NF1. We find that membrane-targeted RSK1 does not impede the culturing of PDAC cells *in vitro*, but rather exhibits inhibitory effects on these cells following the genetic or pharmacological abrogation of mutant KRAS. This suggests that RSK/NF1 participates as a dual negative feedback inhibitor of wild-type KRAS in normal cells (Fig 5E), consistent with previously described RSK1-mediated negative feedback that involves NF1 [59]. The resistance of PDAC cells to membrane-targeted RSK1 may reflect the indifference of oncogenic KRAS to NF1 GAP activity.

### Summary

While prior BirA-RAS studies have included other Ras isoforms and cell types, our experiments more comprehensively focused on PDAC to provide more robust findings in this malignancy [47, 48]. In addition, the normalization of peptide counts to the detection of BirA increased the robustness of the analysis as suggested by the detection of internal and positive controls. It is notable that some biotinylated proteins, such as p110α, could not be verified by MS, suggesting that the sensitivity of MS may be below the threshold for certain proteins and therefore additional fractionation methods should be considered for future BioID-MS work. Nonetheless, our approaches nominated 32 candidate interactors of KRAS^G12D^, 11 of which have been previously reported. Our work showed that ARAF differs from the other two paralogs by being predominantly membrane-localized with mutant RAS compared to the other Raf family members. Since membrane-bound ARAF is transforming even in the absence of active KRAS, this prompts further investigation of potential unique functional roles of ARAF. Other notable mutant- and membrane-specific interactors include the mTORC2 complex described by Kovalski et al. and RSK1, of which we focused on the latter [48].

While RSK1 is known to be involved in the Ras pathway downstream of MAPK1/2 [62], RSK1 does not have an annotated RAS Binding Domain. RSK has been previously described to have negative feedback effects on the Ras pathway via both inhibition of SOS1 and activation of NF1 in the context of wild-type Ras [59, 63]. The labeling of SOS1 by B-G12D was not detected in our study so the inhibition of wild-type KRAS by RSK1 may be predominantly though NF1 activation rather than SOS1 inhibition as reported by Hennig et al. [59]. Although RSK1 and NF1 have been previously described to work in parallel to feedback-inhibit wild-type Ras, our findings demonstrate, for the first time, interaction between NF1 and RSK1. Here, we present two orthogonal approaches to suggest that RSK1 interacts with NF1. In a novel approach combining BioID with CRISPR deletion, we demonstrate that the RSK1 interaction with KRAS^G12D^ is dependent on NF1 expression and SPRED2. This combined with NF1 enrichment by an exogenous RSK1 IP suggest the existence of a protein complex composed of KRAS^G12D^, NF1, RSK1 and SPRED2. Experiments with the MEK inhibitor trametinib further show that the proximity of RSK1 to KRAS^G12D^ is dependent on MAPK signaling. This is consistent with previous studies indicating that RSK1 activity is downstream of MAPK and suggests that the RSK1-KRAS^G12D^ interaction results from the dissociation of RSK1 from MAPK [62, 70]. RSK1 has been previously shown to transiently locate to the membrane upon EGF stimulation and its interaction with NF1 and KRAS may mechanistically explain this observation [61]. We show that membrane targeting of RSK1 does not impair the proliferation of KRAS-mutant PDAC cells; however, membrane-localized RSK inhibits cell proliferation in B-WT and B-G12D/C185S expressing FPC cells upon endogenous mutant KRAS abrogation, which is consistent with RSK1 acting as a negative regulator of Ras signaling.

While we demonstrate that RSK1 attenuates Ras signaling which is subject to negative feedback, the role of this event remains to be determined in KRAS mutant cells. Interestingly, a combinatorial siRNA depletion screen by Yuan et al., suggested that KRAS-mutant cells can be subtyped as either Ras-dependent or RSK-dependent, and PDAC cells were largely the former rather than the latter [71]. Based on our experiments with trametinib in murine PDAC cells, the RSK1/NF1/KRAS^G12D^ complex is dependent on MAPK signaling, which is consistent with MAPK pathway addiction characteristic of the Ras-dependent subtype. On the other hand, RSK-dependent cells are proposed to have various alternate paths to activate RSK and the KRAS^G12D^/NF1/RSK1 complex may prolong the duration of RSK1 at the membrane to achieve events that support oncogenesis. One possibility is that KRAS^G12D^ poises the RSK1/NF1 complex to attenuate wild-type KRAS, which has been purported to be a tumor suppressor [37, 38, 40, 72]. It has been shown that introduction of wild-type KRAS impairs homodimerization of mutant KRAS by forming heterodimers that potentially compete for effectors and membrane docking sites. This feedback-driven mechanism may explain the antagonism between mutant and wild-type KRAS. Another possibility is that given membrane-targeted RSK1 is constitutively active, RSK1 with prolonged residence at the membrane may have greater opportunity to phosphorylate membrane-localized substrates that promote cellular functions associated with tumorigenesis, including increased cell proliferation, survival, migration and glycolytic flux [73–78]. This would implicate a new function for the NF1 tumor suppressor protein in Ras-mutant cells. Consistent with such a function is that biallelic mutations and complete suppression of NF1 is rare in cancer [79]. This motivates further study of the role of NF1 in KRAS-driven PDAC.

## Materials and Methods

### Constructs

The BirA R118G biotin-protein ligase gene including the multiple cloning site was cloned from the pcDNA3.1 mycBioID plasmid deposited by the Roux lab. Murine Kras was inserted between the multiple cloning site of BirA R118G and SalI with GGCGGAAGCGGA, encoding for a short glycine linker (GGSG). The C185S mutation was made with the Q5 Site-Directed Mutagenesis Kit. BirA-CAAX control was created by fusing the last 20 amino acids of murine Kras4B to the C terminus of BirA. The constructs were moved to pBABE-neo between BamHI and SalI for retroviral production. For RSK1 overexpression, human RSK1 cDNA was obtained from John Blenis via Addgene. The sequence (gggagtagcaagagcaagcctaaggaccccagccagcgc) was added after the start codon of the RSK1 cDNA to the fuse a Myr-tag to the N-terminus. Both RSK1 and myr-RSK1 were cloned into pBABE-neo between EcoRI and SalI. Guides were designed using the Benching CRISPR tool against the first coding exon of each target gene with the best off-target scores inserted into the ipUSEPR lentiviral vector [Table S4][171].

### Infections

Phoenix cells and 293T cells were transfected with the Roche X-tremeGENE 9 standard protocol for retrovirus and lentivirus production, respectively, Infected PDAC cells were selected 48 hours post-infection for 5 day if under neomycin selection marker or 2 days if under puromycin selection.

### Cell Culture

Phoenix, 293T, 3T3, mT42, mT93, mT95, and SUIT2 cells were cultured in DMEM with 10% FBS at 37C in 5% CO2 incubator. MiaPACA2 cells were cultured in RPMI with 10% FBS. Cells for BioID experiments had their media changed to serum-free DMEM with 50 μM biotin for 24 hours prior to cell harvest. Cells were washed 5 times with ice cold PBS prior to lysis with mild buffer (50mM HEPES, 150 mM NaCl, 0.7% NP40, 10% Glycerol, 1mM EDTA) with Roche cOmplete Mini protease inhibitor tablet and Roche PhoSTOP phosphatase inhibitor. DC protein assay (Bio-Rad) was used to quantify the protein concentration. 2 μM of 4-hydroxytamoxifen (4-OHT) with DMSO as the vehicle was added to the media for 3 days in FPC experiments. AMG510 (Selleckchem, #S8830) dissolved in DMSO was added to RPMI for MiaPACA2 experiments

### Pulldown and immunoprecipitation

Cell lysates were harvested in mild buffer (50mM HEPES, 150 mM NaCl, 0.7% NP40, 10% Glycerol, 1 mM EDTA) with Roche cOmplete Mini protease inhibitor tablet and Roche PhoSTOP phosphatase inhibitor, at 0 C°. 5mg of lysate was incubated with 30ul of MyOne Streptavidin C1 Dynabeads or 5mg of Myc-Tag (9B11) Mouse mAb (Cell Signaling, #2276) antibody for 3 hours with subsequent 1-hour incubation with 30ul of Protein G Dynabeads for streptavidin pulldown and myc tag IP, respectively. The beads were washed 5 times in lysis buffer before elution in NuPAGE MOPS SDS running buffer.

### Immunoblotting

Streptavidin pulldowns and immunoprecipitations along with their corresponding input whole cell lysates extracts were resolved by NuPAGE 4-12% Bis-Tris gel and transferred on to Immobilon-P PVDF membranes. Membranes were blocked for 1h in 3% BSA/1xTBS/0.1% Tween-20 at RT and incubated overnight with antibodies at 4 C° followed by HRP conjugated secondary antibodies for 30 minutes at RT with 3 TBST washes. Films were exposed to membranes activated with Enhanced Chemiluminescence (ECL) solution.

### Tryptic Digestion

The dynabeads used for SA pulldown were reconstituted with 20uL of 50mM triethylammonium bicarbonate buffer (TEAB). RapiGest was added to a final concentration of 0.1% and tris(2-carboxyethyl) phosphine (TCEP) was added to final concentration of 5mM. Samples were then heated to 55°C for 20min, allowed to cool to room temperature and carboxy ethylmethanethiosulfonate (CEMTS) was added to a final concentration of 10mM. Samples were incubated at room temperature for 20 min to complete blocking of free sulfhydryl groups. 2ug of sequencing grade trypsin (Promega) was then added to the samples and they were digested overnight at 37°C. After digestion the supernatant was removed from the beads and was dried in vacuo. Peptides were reconstituted in 20 ul of 2% acetonitrile: 0.1% formic acid for injection onto the mass spectrometer.

### Mass spectrometry

An Orbitrap Fusion Lumos mass spectrometer (Thermo Scientific), equipped with a nano-ion spray source was coupled to an EASY-nLC 1200 system (Thermo Scientific). The LC system was configured with a self-pack PicoFrit™ 75-μm analytical column with an 8-μm emitter (New Objective, Woburn, MA) packed to 25cm with ReproSil-Pur C18-AQ, 1.9uM material (Dr. Maish GmbH). Mobile phase A consisted of 2% acetonitrile; 0.1% formic acid and mobile phase B consisted of 90% acetonitrile; 0.1% formic Acid. Peptides were then separated using the following steps: at a flow rate of 200 nL/min: 2% B to 6% B over 1 min, 6% B to 30% B over 84 min, 30% B to 60% B over 9 min, 60% B to 90% B over 1 min, held at 90% B for 5 min, 90% B to 50% B over 1 min and then flow rate was increased to 500 nL/min as 50% B was held for 9 min. Eluted peptides were directly electrosprayed into the Fusion Lumos mass spectrometer with the application of a distal 2.3 kV spray voltage and a capillary temperature of 300°C. Full-scan mass spectrum (Res=60,000; 400-1600 m/z) were followed by MS/MS using the “Top Speed” method for selection. High-energy collisional dissociation (HCD) was used with the normalized collision energy set to 35 for fragmentation, the isolation width set to 1.2 and a duration of 10 seconds was set for the dynamic exclusion with an exclusion mass width of 10ppm. We used monoisotopic precursor selection for charge states 2+ and greater, and all data were acquired in profile mode.

### Database Searching

Peaklist files were generated by Proteome Discoverer version 2.2.0.388 (Thermo Scientific). Protein identification was carried out using both Sequest HT and Mascot against the UniProt mouse sequence database (57,220 sequences; 26,386,881 residues) and a database containing Bifunctional ligase/repressor BirA (Escherichia coli). Carboxy ethylthiolation of cysteine was set as fixed modifications, methionine oxidation and n-terminal protein acetylation were set as variable modifications. Trypsin was used as cleavage enzyme with one missed cleavage allowed. Mass tolerance was set at 20 ppm for intact peptide mass and 0.3 Da for fragmentations. Search results were rescored to give a final 1% FDR using a randomized version of the same Uniprot mouse database. Label free quantification was performed by calculating the precursor area for peptides with fit both the Unique and Razor categories in Proteome Discoverer with no imputation and a maximum ratio of 100 between comparison samples. [80, 81]

### Mass Spectrometry Analysis

Proteins quantified by only one peptide were omitted from further analysis. All quantification of peptides was normalized to that of BirA prior to producing relative enrichment by ratiometric analysis. The proteins were ranked by relative enrichment for each biological replicate. The sum of the three assigned ranks was used to indicate a general enrichment score for all three biological replicates to account for protein expression differences across cell lines. The candidate interactors would then be sorted by their cumulative rank scores to identify more consistently enriched interactors. Candidate interactors met their respective criteria if they were enriched in 2 of 3 biological replicates.

### Immunofluorescence

mT93 cells expressing myc tagged constructs were plated on chamber slides for one day prior to being washed with ice cold PBS, fixed with ice cold 4% paraformaldehyde (16% solution, 15710 electron microscopy sciences, diluted with PBS 1:4) for 10 minutes, and quenched with 125μM glycine in PBS three times before permeabilization with 0.5% Triton-X in PBS for 5 minutes. Permeabilization was followed by 3 washes of PBS before blocking in 5% BSA in PBST (0.1% Tween-20) for 1 hour. The myc tag antibody (Cell Signaling, #2276) was added to blocking buffer at a dilution of 1:200 and added to the cells for 2 hours. This incubation was followed by 3 washes prior to secondary antibody incubation with Alexa fluor 568-anti-mouse and Alexa Fluor 488-streptavidin. The cells were followed by 3 washes prior to mounting with ProLong Gold XL antifade mountant with DAPI. The cells were imaged on a Leica SP8 or Zeiss LSM 710 with a 40X objective lens.

### Cell proliferation Assay

Incucyte was used to monitor cell proliferation in SUIT2 cells infected to express both the guide and RFP. CellTiter-Glo Assay was used as instructed in MiaPACA2 cells at days 1,3, and 10. In short, 100 ul of the buffer/substrate mixture was added to each cell-containing well of a 96 well plate and incubated for 10 minutes with gentle shaking prior to being read on a Spectramax i3 plate reader. For crystal violet staining, cells were plated in 6 well plates at a density of 5,000 cells per well and allowed to grow for two days. The cells were washed with ice cold PBS twice before methanol fixation for 20 minutes. These cells were then stained with a crystal violet solution (0.5% crystal violet, 95% ethanol in water).

### 3T3 soft agar Assay

DMEM was supplemented with 10% FBS and 1% Penicillin/Streptomycin. NIH-3T3 cells were infected to express KRAS and BirA constructs and were plated on a bed composed of 1% (w/v) Agar/DMEM in a layer of 0.5% (w/v) Agar/DMEM. After 1 week, the plates were stained with crystal violet solution (0.5% w/v crystal violet, 1% methanol, 1% formaldehyde in PBS), washed and scanned. The foci were manually counted.

## Author Contributions

Category 1

Conception and design of study DK Cheng; DA Tuveson

acquisition of data: DK Cheng; N Prasad; KD Rivera;

analysis and/or interpretation of data: DK Cheng; DJ Pappin; DA Tuveson

Category 2

Drafting the manuscript: DK Cheng; TE Oni; DA Tuveson

revising the manuscript critically for important intellectual content: DK Cheng; TE Oni, JS

Thalappillil; B Alagesan; Y Park; DA Tuveson

Category 3

Approval of the version of the manuscript to be published: DK Cheng; TE Oni; B Alagesan; Y Park; JS Thalappillil; N Prasad; KD Rivera; DJ Pappin; DA Tuveson

## Acknowledgments

We thank Dr. John Blenis and Dr. Joe W. Ramos and Dr. Channing Der for advice. We acknowledge the Cold Spring Harbor Laboratory Microscopy and Mass Spectrometry Shared Resources, which are supported by the NIH Cancer Center Support Grant P30CA045508. We thank members of the Tuveson Laboratory for their assistance and advice. DAT is a distinguished scholar of the Lustgarten Foundation and Director of the Lustgarten Foundation-designated Laboratory of Pancreatic Cancer Research. DAT is also supported by the Cold Spring Harbor Laboratory Association, the V Foundation, the Thompson Foundation, and the National Institutes of Health (NIH P30CA45508, P20CA192996, U10CA180944, U01CA224013, U01CA210240, R01CA188134, and R01CA190092). DAT is supported by the Simons Foundation (552716). YP is supported by NCI R50CA211506. BA is supported by NCI F30CA200240. DC is supported by NCI F30CA213883. JST is supported by NCI F31CA247416. This work was also supported by the Cold Spring Harbor Laboratory and Northwell Health Affiliation (Project Lazarus).

## Supporting information

**Fig S1A.**
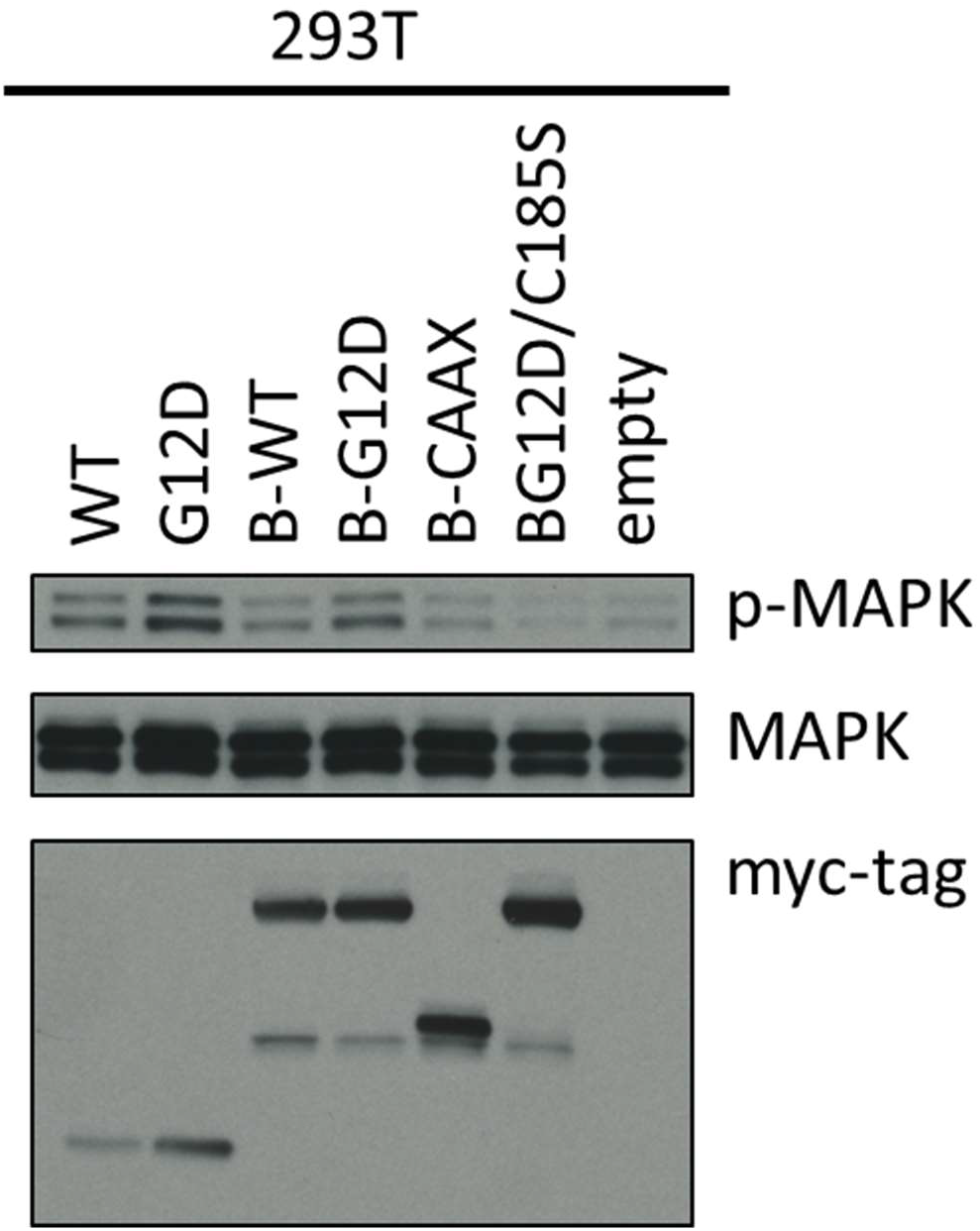
BirA-KrasG12D induces MAPK phosphorylation in 293T cells. 293T cells expressing KRAS and BirA-KRAS fusions were harvested for western blot analysis of MAPK activation.

**Fig S1B.**
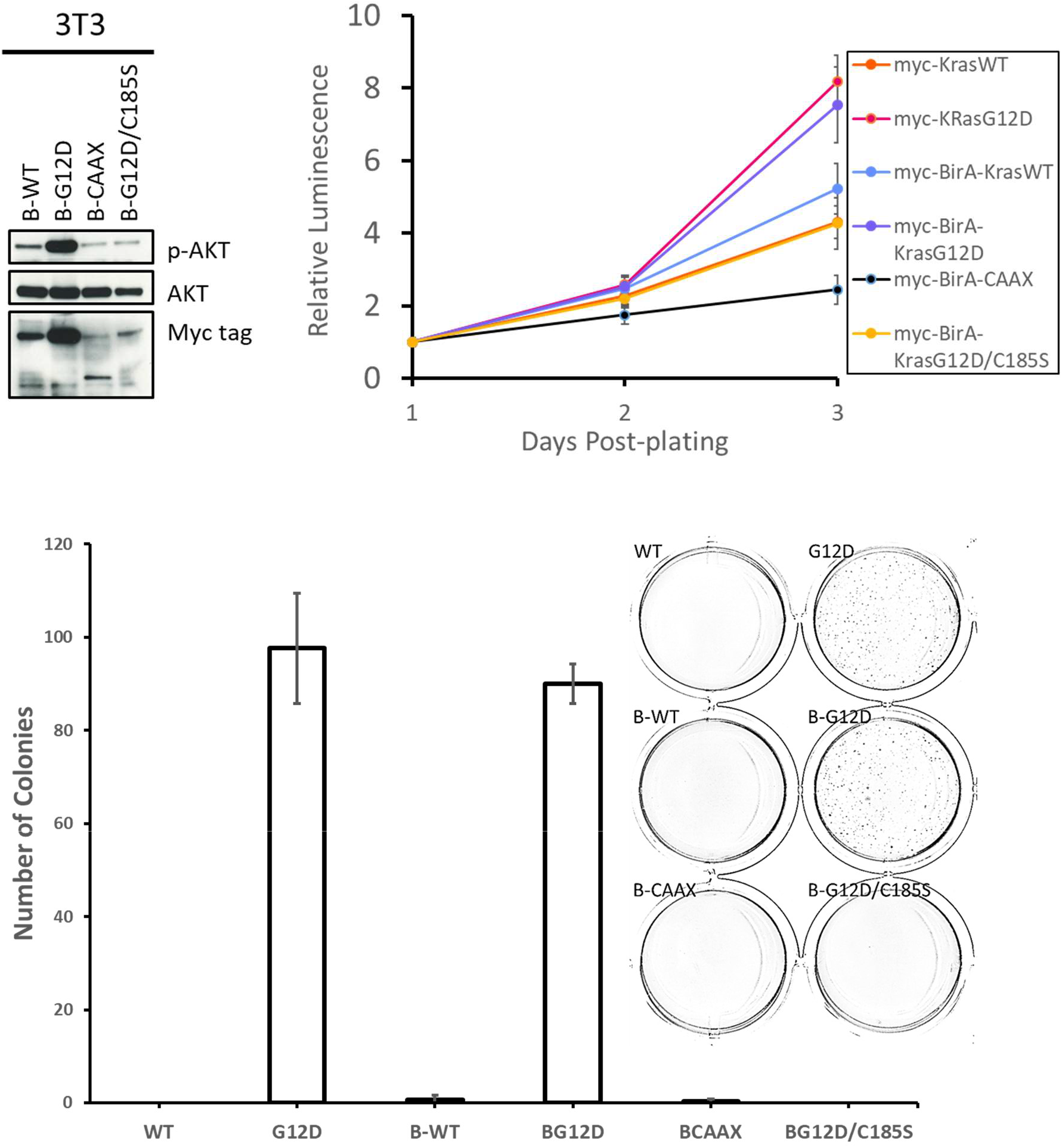
BirA-KRAS^G12D^ expression functionally phenocopies KRAS^G12D^ expression in 3T3 cells with regards to cell proliferation and anchorage independent growth. Infection of 3T3 cells to express BirA fusion proteins along with myc-tagged KRAS controls was performed and selected for 5 days followed by Western blot analysis of expression of BirA constructs along with phosphorylation status of AKT (top left). Proliferation assays (top right) for 3T3 cells expressing myc-tagged KRAS and BirA KRAS fusion proteins. Relative luminescent was normalized to Day1 and measured every 24 hours for three days. Error bars represent standard error of the mean (n=5) and soft agar assays (bottom) for 3T3 cells. Number of colonies were counted in each well. Error bars represent standard error of the mean of technical replicates (n=2)

**Fig S2.**
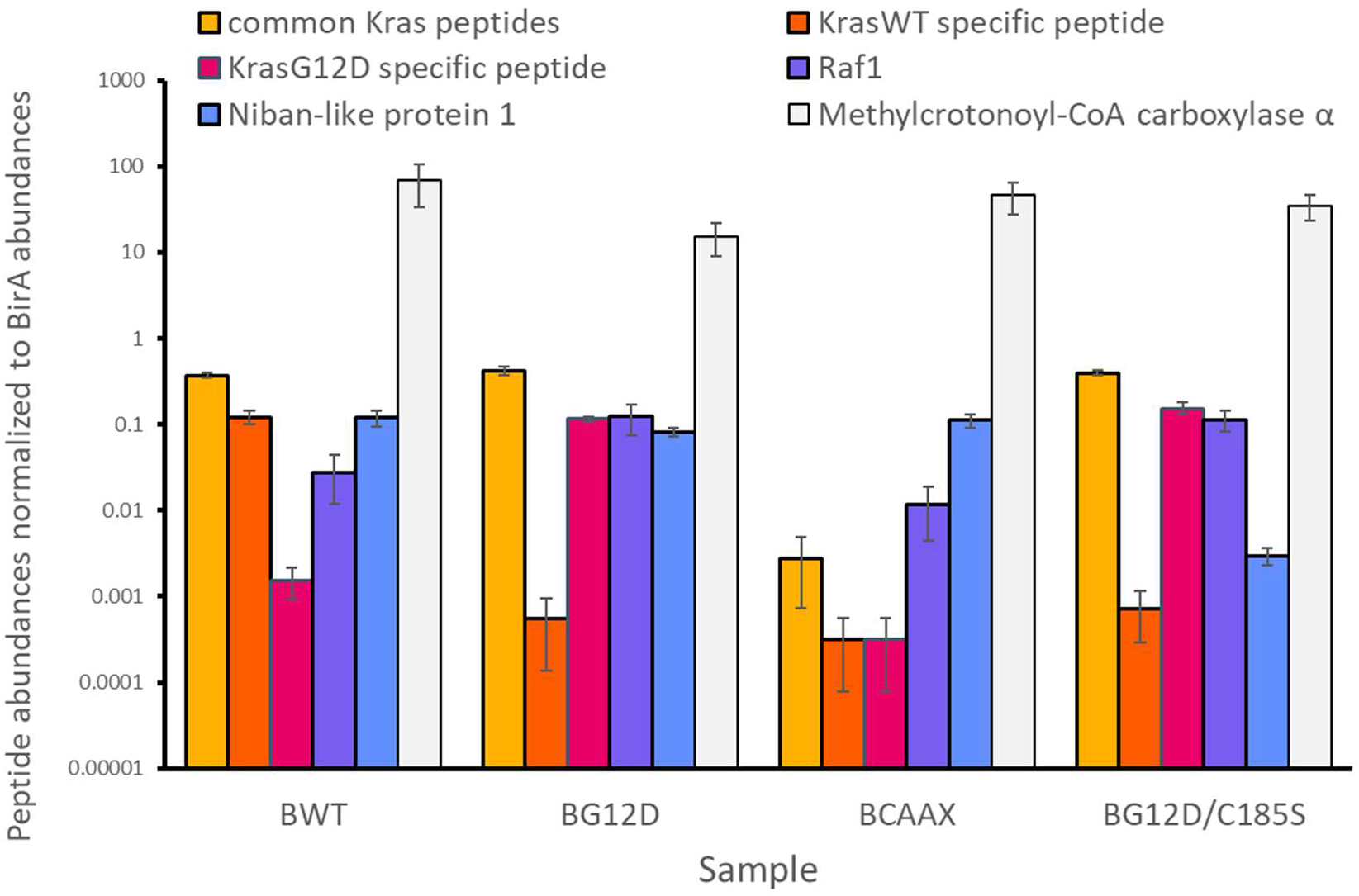
Mass spectrometry identifies internal and positive control peptides. Mass spectrometry relative quantification of internal control KRAS peptides and positive controls (RAF1 and niban-like 1) after normalization to BirA counts for B-WT, B-G12D, B-CAAX and B-G12D/C185S samples from 4-OHT treated FPC cells. Error bars represent standard error of the mean (n=3)

**Fig S3A.**
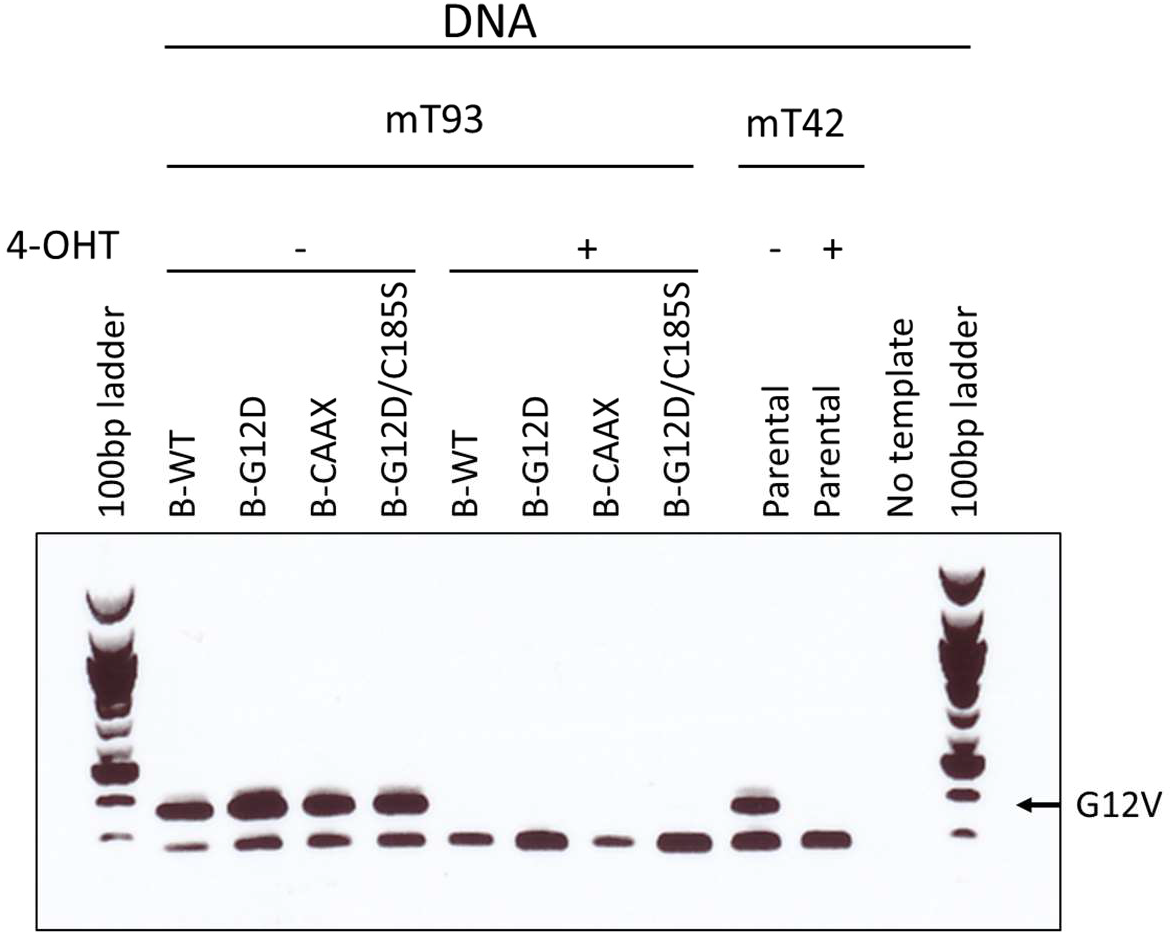
4-hydroxy-tamoxifen induces the excision of Kras^G12V^ allele. mT93 FPC cells expressing the panel of BirA-Kras fusion proteins were treated with either vehicle or 4-OHT for 3 days. genomic DNA was isolated and amplified for the *Kras^G12V^* allele.

**Fig S3B.**
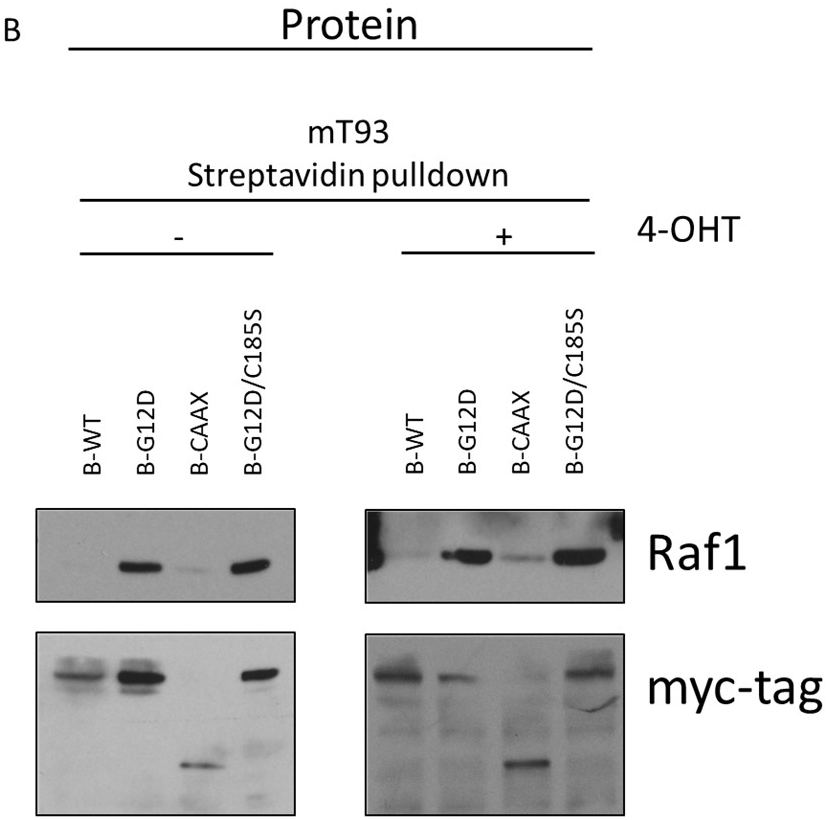
Labelling of Raf1 by the BioID-KRAS assay on cells with and without endogenous mutant Kras expression.

**Fig S3C.**
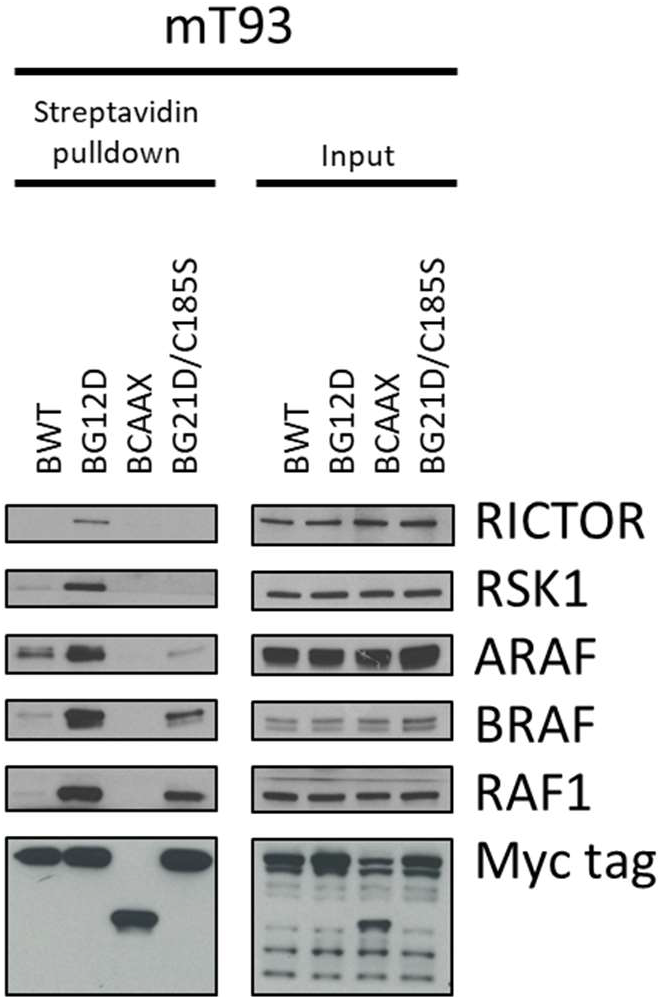
Immunoblot confirmation of biotinylation of RICTOR, RSK1, ARAF, BRAF, and RAF1 as B-G12D substrates in mT93.

**Fig S3D.**
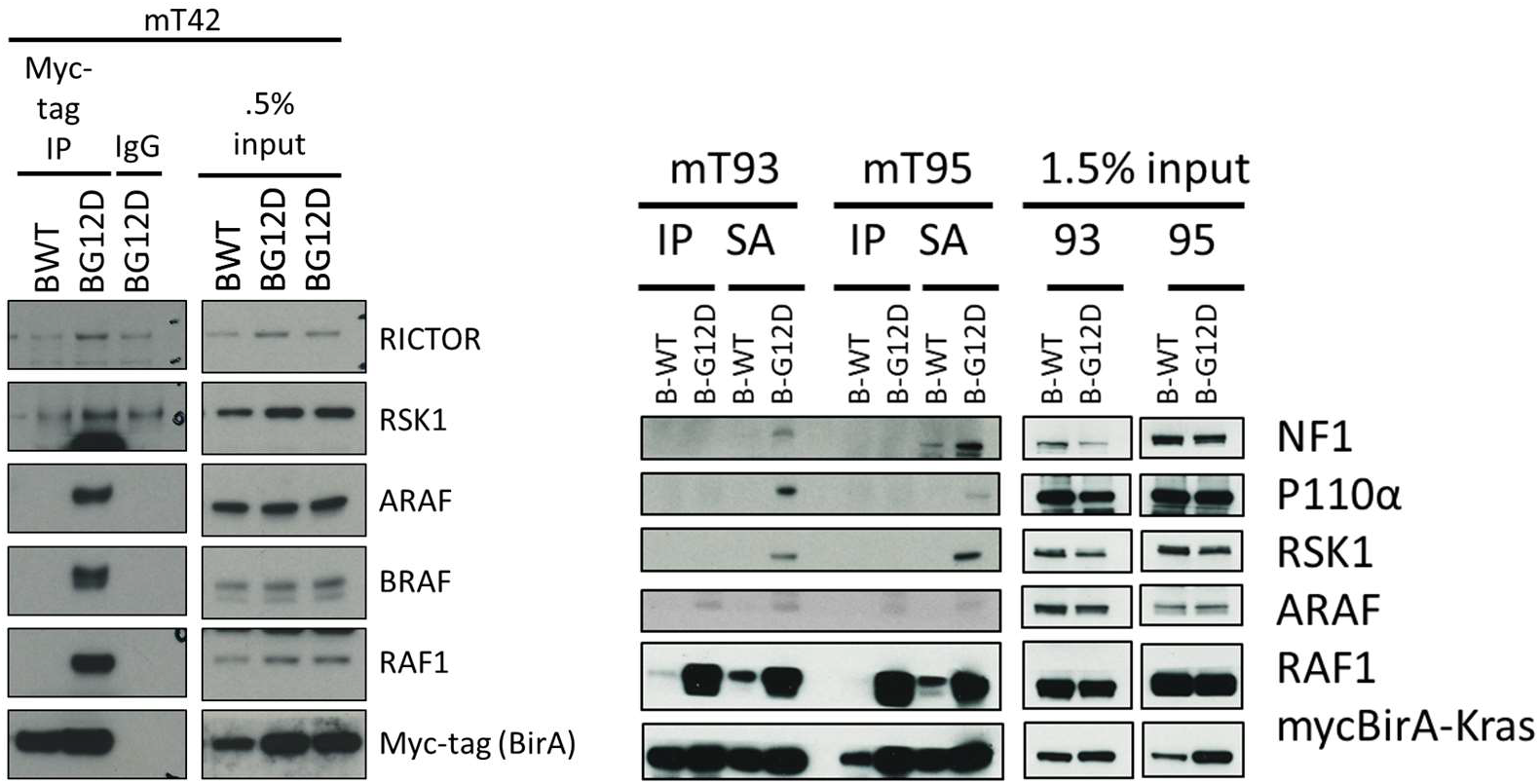
Comparison of immunoprecipitation to BioID in PDAC cell lines. (Left) Lysates of B-WT and B-G12D expressing mT42 cells incubated with beads coated with myc-tag antibodies for IP. Proteins were eluted for western blot analysis. (Right) Lysates of B-WT and B-G12D-expressing mT93 and mT95 cells were used as input for both immunoprecipitation and streptavidin pulldown.

**Fig S4.**
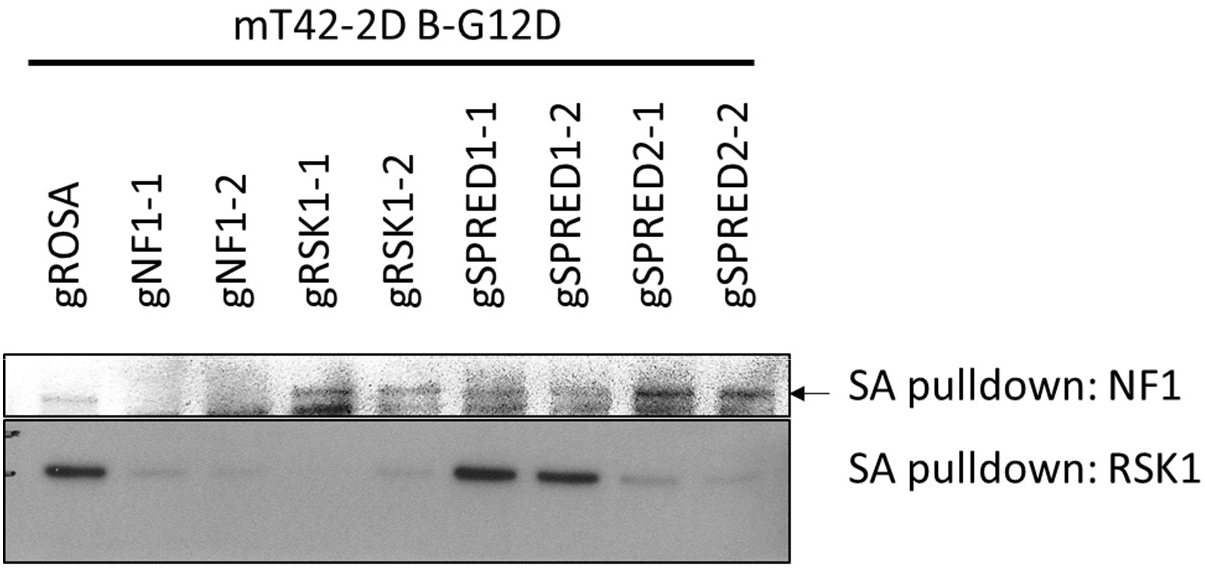
The BioID assay was performed on mT42 cells expressing B-G12D and guides against NF1, RSK1, SPRED1, and SPRED2.

**Fig S5A.**
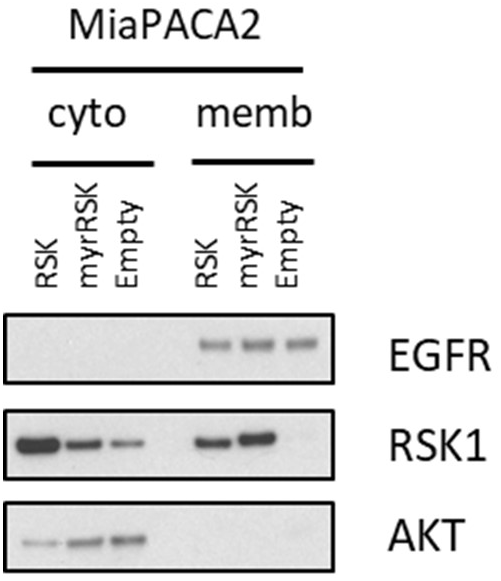
Myr-tagged RSK preferentially localizes to the plasma membrane compartment. MiaPACA2 cells were infected to express RSK and myr-tagged RSK1. Proteins from cytosolic and membrane compartments were extracted with a sequential fractionation protocol [54].

**Fig S5B.**
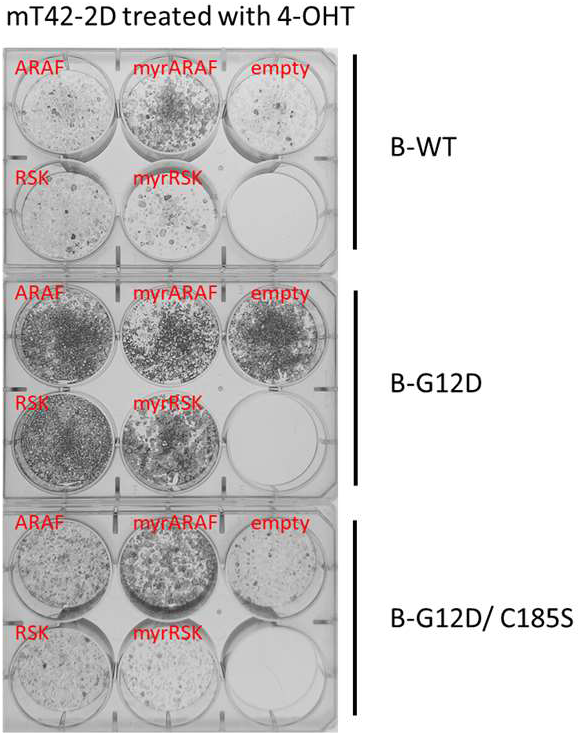
Myr-RSK further inhibits murine PDAC cells upon endogenous Kras^G12V^ excision. mT42 cells expressing B-WT, B-G12D, B-G12D/C185S were infected to express ARAF, myr-tagged ARAF, RSK1, myr-tagged RSK1, and empty control. After selection, the cells were sparsely plated at 5000 cells per well with 4-OHT and allowed to grow for 7 days. The cells were stained with crystal violet staining. Crystal violet was dissolved with ethanol and quantified.

**Fig S5C.**
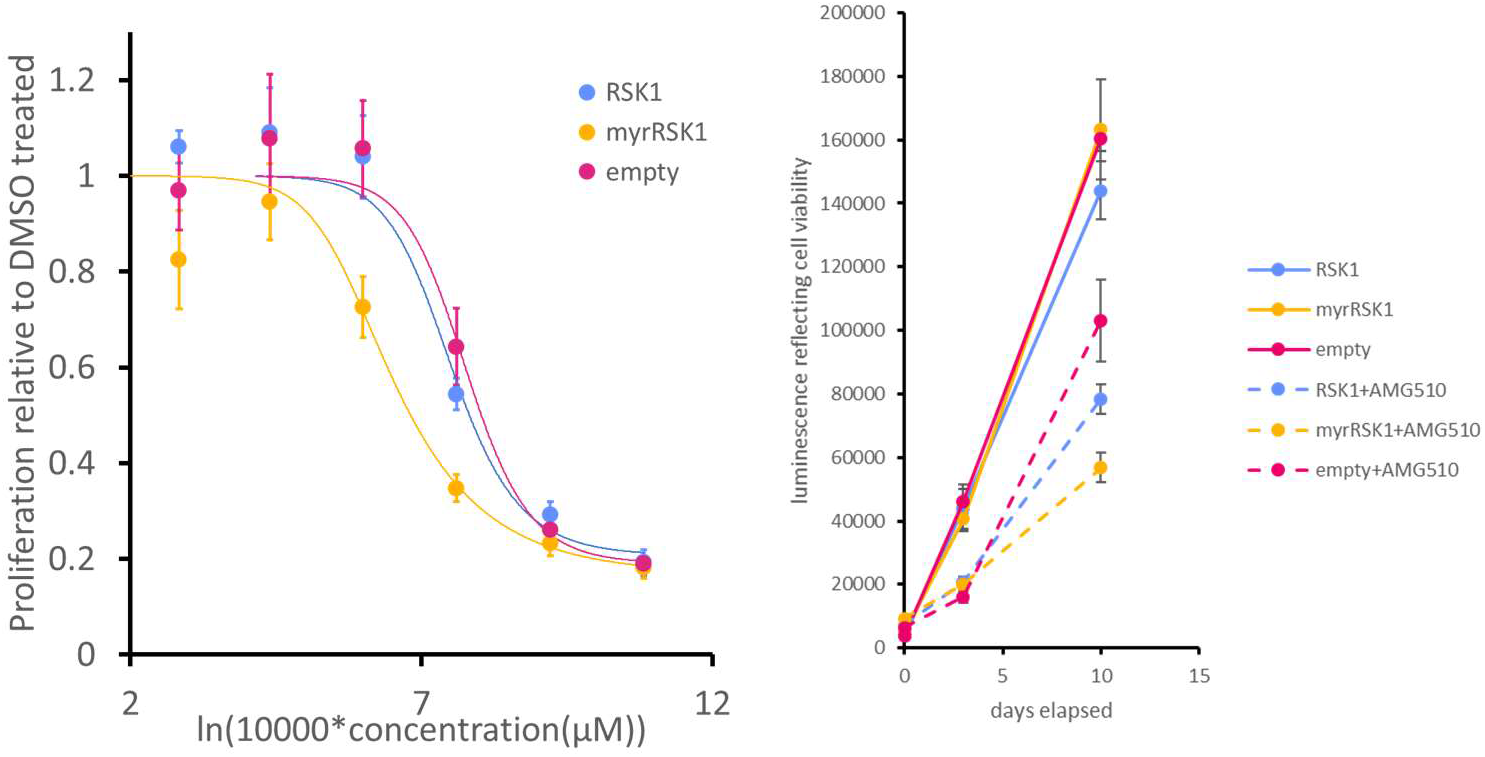
myr-tagged RSK1 sensitizes MiaPACA2 cells to KRAS^G12C^ inhibition. (Left) MiaPACA2 cells expressing RSK1, myr-RSK1, and empty control were treated with various concentrations of AMG510 (5 μM, 1 μM, 200 nM, 40nM, 8nM, 1.6nM). Proliferation was measured by relative confluency by Incucyte imaging. (Right) cell viability assays were performed at day of plating with 3 and 10 days post-plating on MiaPACA2 cells expressing RSK1, myr-RSK1, and empty comtrol were treated with 200 nM of AMG510. Error bars represent standard deviation of technical replicate wells.

**Fig S5D.**
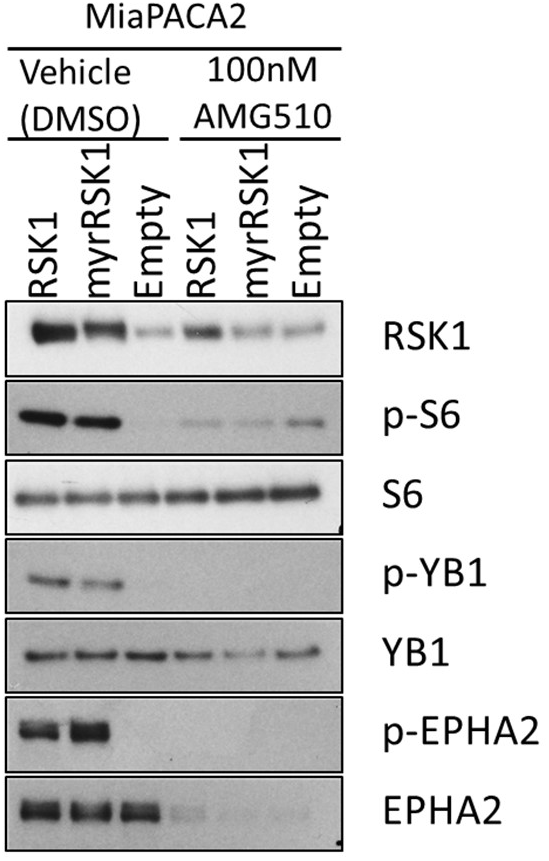
myr-RSK expresion is unstable in MiaPACA2 cells upon KrasG12C inhibition. MiaPACA2 cells expressing RSK1 and myrRSK1 along with infected control were treated with AMG510 or DMSO control over 2 days. Harvested lysates were probed for RSK expression and phosphorylation status of RSK substrates (S6, YB-1 and EPHA2)

**Table S1.**
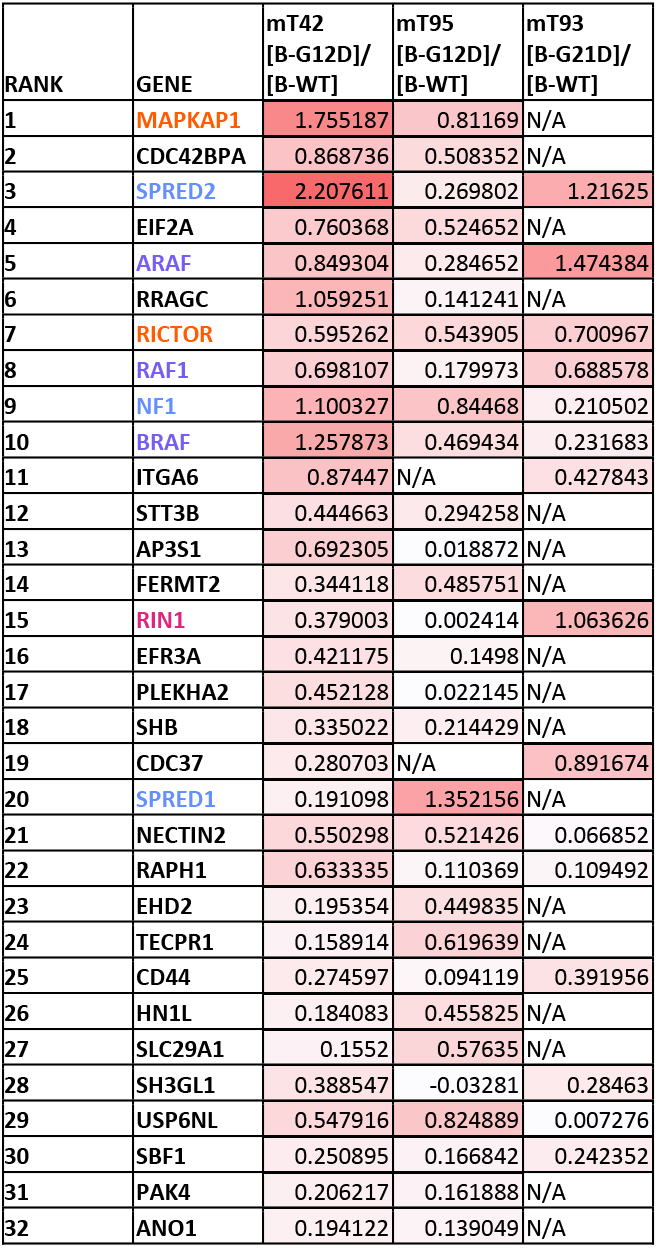
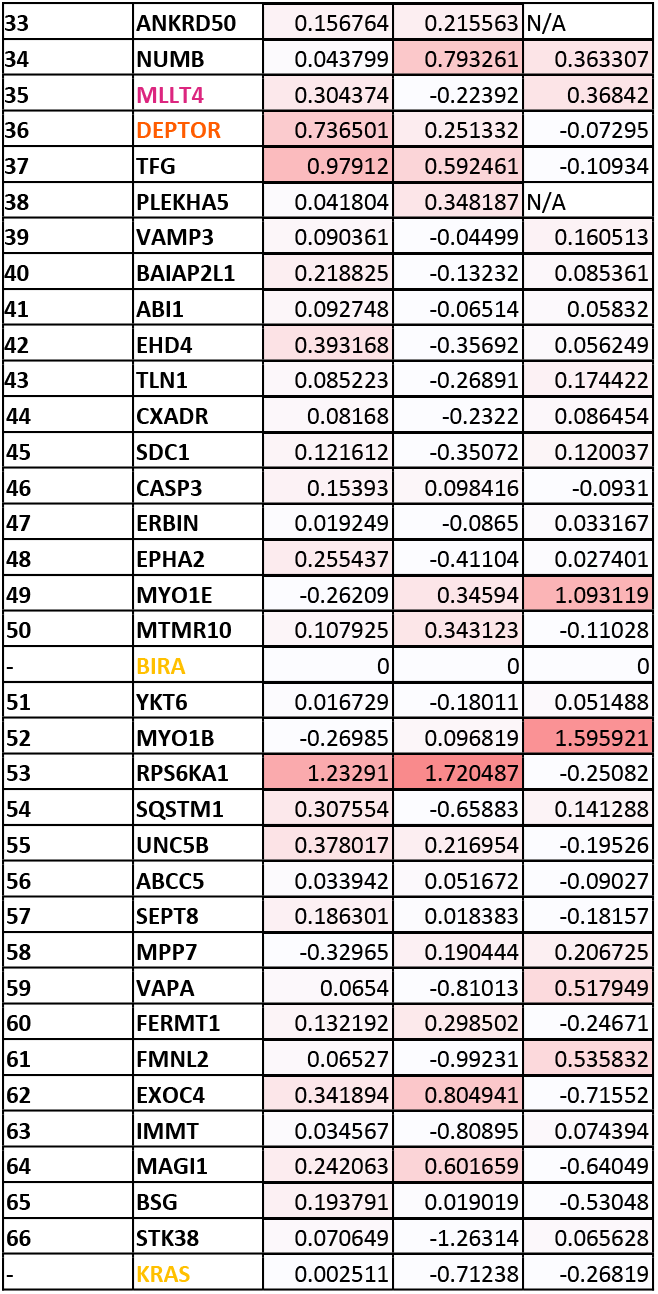
66 Candidate mutant specific interactors were nominated based on enrichment in B-G12D compared to B-WT. Log_10_ enrichment of candidate interactor peptides of B-G12D compared to B-WT after normalization to BirA counts. Known KRAS interactors include RAF members (purple), NF1/SPRED proteins (blue), mTORC2 (orange), and RBD-containing proteins (magenta) with the internal controls, BirA and KRAS (yellow).

**Table S2.** ARAF is preferentially biotinylated by B-G12D compared to B-G12D/C185S. Log_10_ enrichment of B-G12D compared to B-G12D/C185S after normalization to BirA counts for candidate interactor peptides. Known KRAS interactors include RAF members (purple), NF1/SPRED proteins (blue), mTORC2 (orange), and RBD-containing proteins (magenta) with the internal controls, BirA and KRAS (yellow).

| RANK | GENE | mT42<br>[B-G12D]/<br>[B-G12D/C185S] | mT95<br>[B-G12D]/<br>[B-G12D/C185S] | mT93<br>[B-G12D]/<br>[B-G12D/C185S] |
| --- | --- | --- | --- | --- |
| 1 | CXADR | 2.77335 | 2.391534 | 0.641675 |
| 2 | MAPKAP1 | 1.964699 | 1.571517 | N/A |
| 3 | SEPT8 | 1.638286 | 1.7218 | 0.901369 |
| 4 | CD44 | 2.318534 | 1.129588 | 1.357382 |
| 5 | YKT6 | 3.091127 | 1.020428 | 2.381236 |
| 6 | DEPTOR | 1.879239 | 1.208165 | 0.99095 |
| 7 | NF1 | 1.375172 | 1.277483 | 0.634414 |
| 8 | ABI1 | 1.077095 | 1.98066 | 1.184148 |
| 9 | VAMP3 | 1.703178 | 0.753709 | 0.789873 |
| 10 | PLEKHA5 | 1.726858 | 0.688121 | N/A |
| 11 | USP6NL | 1.191218 | 1.164823 | 0.805178 |
| 12 | MTMR10 | 1.923092 | 0.535914 | 0.693137 |
| 13 | EXOC4 | 1.069929 | 1.144875 | 0.559587 |
| 14 | EFR3A | 1.490905 | 0.618939 | N/A |
| 15 | SHB | 1.618939 | 0.554363 | N/A |
| 16 | BAIAP2L1 | 0.998371 | 0.70001 | 1.180474 |
| 17 | ABCC5 | 1.359178 | 0.391606 | 1.037121 |
| 18 | CDC42BPA | 0.626974 | 1.234939 | N/A |
| 19 | SLC29A1 | 0.600387 | 0.916284 | N/A |
| 20 | UNC5B | 2.119343 | 0.07753 | 1.684728 |
| 21 | FMNL2 | 1.035895 | 0.344671 | 0.959745 |
| 22 | ANKRD50 | 1.411051 | 0.116527 | N/A |
| 23 | ERBIN | 0.907435 | 0.356142 | 0.654053 |
| 24 | EPHA2 | 1.159129 | 0.158101 | 0.505675 |
| 25 | NECTIN2 | 2.40406 | -0.00141 | 1.134362 |
| 26 | ANO1 | 0.680206 | 0.478983 | N/A |
| 27 | EHD2 | 0.404865 | 1.027629 | N/A |
| 28 | SPRED1 | 1.302899 | 0.082412 | N/A |
| 29 | NUMB | 0.470505 | 0.66808 | 1.822717 |
| 30 | TECPR1 | 0.320962 | 0.959573 | N/A |
| 31 | ARAF | 0.553303 | 0.58561 | 1.346112 |
| 32 | FERMT2 | 0.319881 | 0.754161 | N/A |
| 33 | PLEKHA2 | 1.14736 | 0.058294 | N/A |
| 34 | STT3B | 0.986879 | 0.106333 | N/A |
| 35 | RICTOR | 0.485484 | 0.410654 | 1.277203 |
| 36 | AP3S1 | 0.901817 | 0.092125 | N/A |
| 37 | BSG | 1.724652 | 1.727046 | 0.341779 |
| 38 | MPP7 | 0.60966 | 0.198911 | 1.174833 |
| 39 | HN1L | 0.128424 | 0.795759 | N/A |
| 40 | EHD4 | 0.23357 | 0.547979 | 0.588656 |
| 41 | SPRED2 | 0.476836 | 0.204901 | 2.014316 |
| 42 | PAK4 | 0.561277 | 0.13151 | N/A |
| 43 | SBF1 | 2.143627 | 0.683091 | 0.312754 |
| 44 | SH3GL1 | -0.01074 | 1.311871 | 0.464558 |
| 45 | MAGI1 | 0.038351 | 0.941593 | N/A |
| 46 | SDC1 | 0.080284 | 0.539005 | 0.54395 |
| 47 | RRAGC | 0.078709 | 0.481175 | N/A |
| 48 | VAPA | 0.692765 | -0.14713 | 0.941862 |
| 49 | TLN1 | 0.111609 | 0.125661 | 0.938667 |
| 50 | EIF2A | 0.045417 | 0.245924 | N/A |
| 51 | RIN1 | 0.117073 | 0.009992 | 0.52856 |
| 52 | MLLT4 | 0.213892 | 0.160183 | 0.421403 |
| 53 | RPS6KA1 | 1.021264 | 0.626588 | -0.0483 |
| 54 | CASP3 | 0.5738 | -0.08546 | 0.272262 |
| 55 | TFG | -0.01791 | 0.159397 | 0.314574 |
| 56 | BRAF | 0.074277 | -0.22239 | 0.461109 |
| 57 | RAF1 | -0.10195 | 0.133899 | 0.288583 |
| 58 | MYO1B | -0.30977 | 0.099798 | 1.021896 |
| 59 | MYO1E | -0.4131 | 0.163004 | 0.69719 |
| - | BIRA | 0 | 0 | 0 |
| 60 | ITGA6 | 2.124571 | -1.30643 | 0.851755 |
| - | KRAS | -0.02896 | -0.06005 | 0.057823 |
| 61 | RAPH1 | 1.502431 | -0.9417 | 0.231125 |
| 62 | CDC37 | 0.490215 | -0.97811 | 0.374808 |
| 63 | FERMT1 | 0.876572 | -1.26993 | 0.638027 |
| 64 | SQSTM1 | -0.30324 | 0.037976 | 0.101571 |
| 65 | STK38 | 0.542777 | -1.18742 | 0.469576 |
| 66 | IMMT | 0.022742 | -0.60851 | 0.143171 |

**Table S3.** ARAF and RSK1 are top mutant and membrane specific candidate interactors of KRAS^G12D^. Log_10_ enrichment of counts for candidate interactor peptides in the B-G12D sample compared to that of B-WT after normalization to BirA sorted by enrichment following excision of *Kras^G12V^* in FPC cells. Known KRAS interactors include RAF members (purple), NF1/SPRED proteins (blue), mTORC2 (orange), and RBD-containing proteins (magenta) with the internal controls, BirA and KRAS (yellow).

| Rank | Gene Name | mT42 [B-G12D]/[B-WT] | mT95 [B-G12D]/[B-WT] | mT93 [B-G21D]/[B-WT] | mT42 [B-G12D]/[B-WT] +4OHT | mT95 [B-G12D]/[B-WT] +4OHT | mT93 [B-G21D]/[B-WT] +4OHT |
| --- | --- | --- | --- | --- | --- | --- | --- |
| 1 | ARAF | 0.849304 | 0.284652 | 1.474384 | 0.787205 | 0.91922 | 1.455849 |
| 2 | RPS6KA1 | 1.23291 | 1.720487 | -0.25082 | 1.117148 | 0.67013 | 1.108004 |
| 3 | SPRED1 | 0.191098 | 1.352156 | N/A | 0.534964 | 1.143845 | N/A |
| 4 | EFR3A | 0.421175 | 0.1498 | N/A | 1.203169 | 0.574156 | N/A |
| 5 | SPRED2 | 2.207611 | 0.269802 | 1.21625 | 0.201151 | 1.449408 | 1.417899 |
| 6 | RAF1 | 0.698107 | 0.179973 | 0.688578 | 1.033701 | 0.606426 | 0.567235 |
| 7 | CDC42BPA | 0.868736 | 0.508352 | N/A | 0.905865 | 0.456833 | N/A |
| 8 | DEPTOR | 0.736501 | 0.251332 | -0.07295 | 0.663505 | 0.483558 | 0.363096 |
| 9 | MAPKAP1 | 1.755187 | 0.81169 | N/A | 1.509053 | 0.495902 | 0.158092 |
| 10 | NF1 | 1.100327 | 0.84468 | 0.210502 | 1.004831 | 0.18394 | N/A |
| 11 | MYO1B | -0.26985 | 0.096819 | 1.595921 | -0.06986 | 0.973282 | 1.405838 |
| 12 | ITGA6 | 0.87447 | N/A | 0.427843 | 0.621476 | 0.721674 | 0.015789 |
| 13 | RICTOR | 0.595262 | 0.543905 | 0.700967 | 0.477894 | 0.066503 | 0.847274 |
| 14 | PLEKHA2 | 0.452128 | 0.022145 | N/A | 0.649327 | 0.15541 | N/A |
| 15 | USP6NL | 0.547916 | 0.824889 | 0.007276 | 0.412789 | -0.02865 | 0.519207 |
| 16 | RIN1 | 0.379003 | 0.002414 | 1.063626 | 0.343432 | -0.1328 | 1.208626 |
| 17 | MYO1E | -0.26209 | 0.34594 | 1.093119 | -0.11432 | 0.753183 | 0.626953 |
| 18 | PAK4 | 0.206217 | 0.161888 | N/A | 0.340043 | 0.105114 | N/A |
| 19 | EHD2 | 0.195354 | 0.449835 | N/A | 0.036919 | 0.531507 | N/A |
| 20 | MPP7 | -0.32965 | 0.190444 | 0.206725 | -0.15341 | 0.382771 | 0.260023 |
| 21 | SH3GL1 | 0.388547 | -0.03281 | 0.28463 | 0.334933 | 0.561053 | -0.10902 |
| 22 | ABCC5 | 0.033942 | 0.051672 | -0.09027 | 0.313796 | -0.30816 | 0.314398 |
| 23 | MLLT4 | 0.304374 | -0.22392 | 0.36842 | 0.22508 | -0.34028 | 0.448115 |
| 24 | NECTIN2 | 0.550298 | 0.521426 | 0.066852 | 0.04035 | -0.20411 | 0.469357 |
| 25 | CXADR | 0.08168 | -0.2322 | 0.086454 | 0.095287 | -0.08919 | 0.089288 |
| 26 | CASP3 | 0.15393 | 0.098416 | -0.0931 | 0.235395 | -0.41959 | 0.229345 |
| 27 | VAPA | 0.0654 | -0.81013 | 0.517949 | 0.097546 | 0.377074 | -0.10596 |
| - | BIRA | 0 | 0 | 0 | 0 | 0 | 0 |
| - | KRAS | 0.002511 | -0.71238 | -0.26819 | 0.08982 | 0.07801 | -0.0613 |
| 28 | SBF1 | 0.250895 | 0.166842 | 0.242352 | 0.442565 | 0.191822 | -0.20449 |
| 29 | BRAF | 1.257873 | 0.469434 | 0.231683 | 1.207007 | 0.161276 | -0.27541 |
| 30 | VAMP3 | 0.090361 | -0.04499 | 0.160513 | 0.289234 | 0.13625 | -0.26325 |
| 31 | BAIAP2L1 | 0.218825 | -0.13232 | 0.085361 | -0.5608 | 0.07819 | 0.181615 |
| 32 | SQSTM1 | 0.307554 | -0.65883 | 0.141288 | 0.033291 | 0.560629 | -0.31531 |

**Table S4.**
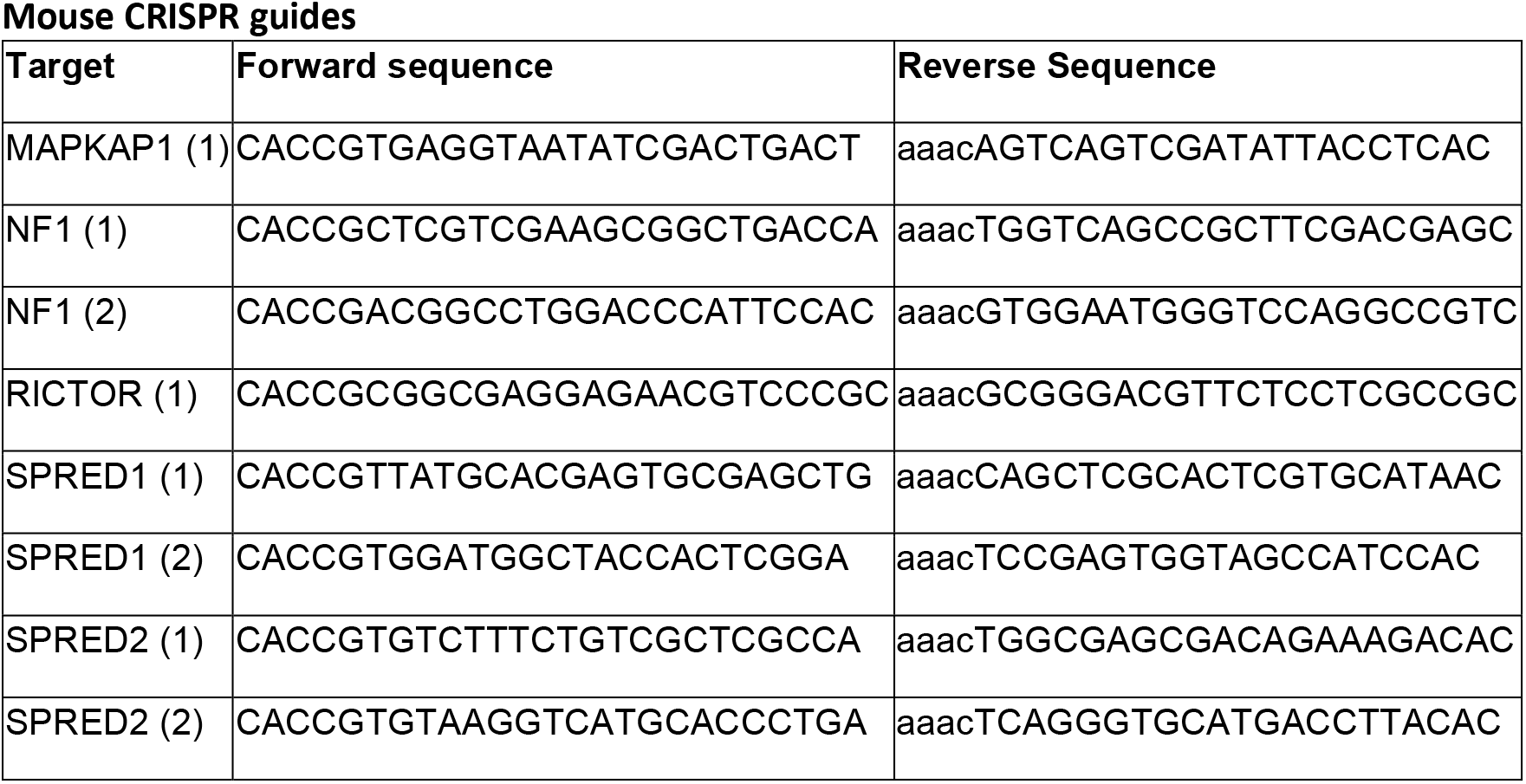
Sequences of Mouse CRISPR guides used. Guides were designed in Benchling and selected based on best off-target scores.

